# A wild *Cucurbita* genome reveals the role of structural variants and introgression in domestication

**DOI:** 10.1101/2020.10.15.341990

**Authors:** Josué Barrera-Redondo, Guillermo Sánchez-de la Vega, Jonás A. Aguirre-Liguori, Gabriela Castellanos-Morales, Yocelyn T. Gutiérrez-Guerrero, Xitlali Aguirre-Dugua, Erika Aguirre-Planter, Maud I. Tenaillon, Rafael Lira-Saade, Luis E. Eguiarte

**Affiliations:** Departamento de Ecología Evolutiva, Instituto de Ecología, Universidad Nacional Autónoma de México, Circuito Exterior s/n Anexo al Jardín Botánico, 04510 Ciudad de México, México; Department of Ecology and Evolutionary Biology, University of California, Irvine, CA; Departamento de Conservación de la Biodiversidad, El Colegio de la Frontera Sur, Villahermosa, Carretera Villahermosa-Reforma km 15.5 Ranchería El Guineo 2ª sección, 86280 Villahermosa, Tabasco, México; Génétique Quantitative et Evolution – Le Moulon, Université Paris-Saclay, Institut National de Recherche pour l’Agriculture, l’Alimentation et l’Environnement, Centre National de la Recherche Scientifique, AgroParisTech, Gif-sur-Yvette, France; UBIPRO, Facultad de Estudios Superiores Iztacala, Universidad Nacional Autónoma de México, Av. de los Barrios #1, Col. Los Reyes Iztacala, Tlalnepantla, Edo. de Mex 54090, México

**Keywords:** Squash, domestication, Mesoamerica, structural variants, selective scans

## Abstract

Despite their economic importance and well-characterized domestication syndrome, the genomic impact of domestication and the identification of variants underlying the domestication traits in *Cucurbita* species (pumpkins and squashes) is currently lacking. *Cucurbita argyrosperma*, also known as cushaw pumpkin or silver-seed gourd, is a Mexican crop consumed primarily for its seeds rather than fruit flesh. This makes it a good model to study *Cucurbita* domestication, as seeds were an essential component of early Mesoamerican diet and likely the first targets of human-guided selection in pumpkins and squashes. We obtained population-level data using tunable Genotype by Sequencing libraries for 192 individuals of the wild and domesticated subspecies of *C. argyrosperma* across Mexico. We also assembled the first wild *Cucurbita* genome at a chromosome level. Comparative genomic analyses revealed several structural variants and presence/absence of genes related to domestication. Our results indicate a monophyletic origin of this domesticated crop in the lowlands of Jalisco. We uncovered candidate domestication genes that are involved in the synthesis and regulation of growth hormones, plant defense mechanisms, flowering time and seed development. The presence of shared selected alleles with the closely related species *Cucurbita moschata* suggests domestication-related introgression between both taxa.

## Introduction

Domestication is an evolutionary process where human societies select, modify and eventually assume control over the reproduction of useful organisms. A mutualistic relationship emerges from this interaction, where humans exploit a particular resource of interest, while the domesticated organism benefits from increased fitness and extended geographical range (Meyer and Purugganan, 2013; Zeder, 2015). This is well illustrated in *Cucurbita* L. (pumpkins, squashes and some gourds), where human-guided domestication and breeding have considerably extended their distribution despite the extinction of their natural dispersers (*e.g.*, mastodons and similar megafauna) (Kistler et al., 2015). Today, *Cucurbita* stand as successful crops grown and consumed worldwide, with a global annual production of ~24 million tons (Paris, 2016).

With ca. 21 taxa, the *Cucurbita* genus has experienced independent domestication events in five species (Castellanos-Morales et al., 2018; Sanjur et al., 2002). Each *Cucurbita* crop experienced unique selection for specific traits, predominantly defined by the nutritional and cultural needs of early human populations in America (Zizumbo-Villarreal et al., 2012). However, many domestication traits are common to domesticated *Cucurbita*, including the loss of bitter compounds (cucurbitacins), the loss of physical defense mechanisms (*e.g*., urticating trichomes), the loss of seed dormancy, the enlargement of fruits and seeds, and the diversification of fruit morphology (Chomicki et al., 2019; Paris, 2016).

The initial steps of *Cucurbita* domestication were most likely directed towards seed rather than flesh consumption (Whitaker and Cutler, 1965). Seeds are rich in both carbohydrates and fatty acids, and cucurbitacins can be removed through boiling and washing; processes that are still employed for the consumption of wild *Cucurbita* seeds in Western Mexico (Zizumbo-Villarreal et al., 2012). Because the cultivation of *C. argyrosperma* (pipiana squash, cushaw pumpkin or silver-seed gourd) is directed towards seed production rather than fruit flesh, it stands as an excellent model to investigate the early steps of *Cucurbita* domestication. *Cucurbita argyrosperma* subsp. *argyrosperma* (*argyrosperma* hereafter) was domesticated in Mesoamerica from its wild relative *Cucurbita argyrosperma* subsp. *sororia* (*sororia* hereafter), according to archaeological and genetic evidence (Sanjur et al., 2002; Piperno et al., 2009; Sánchez-de la Vega et al., 2018). *Argyrosperma* exhibits morphological differences from *sororia*, including larger fruits, larger seeds and lack of urticating trichomes (Fig. 1). The earliest archaeological record of *argyrosperma* is presumed to be from 8,700-year-old phytoliths in the Central Balsas Valley (Guerrero), although its taxonomic identity remains uncertain (Piperno et al., 2009). Its earliest unambiguous archaeological remain is a 5,035-year-old peduncle from the Ocampo caves in Tamaulipas (Smith, 1997). *C. argyrosperma* is a monoicous outcrossing species and gene flow has been previously described between the domesticated and wild subspecies (Montes-Hernandez and Eguiarte, 2002). Both subspecies are sympatrically distributed throughout the Pacific Coast of Mexico and Central America, with a few populations scattered in the coast of the Gulf of Mexico (Lira et al., 2016; Sánchez-de la Vega et al., 2018). The domesticated taxon is also distributed in the Yucatan Peninsula, where its wild counterpart is absent (Sánchez-de la Vega et al., 2018).

**Figure 1.**
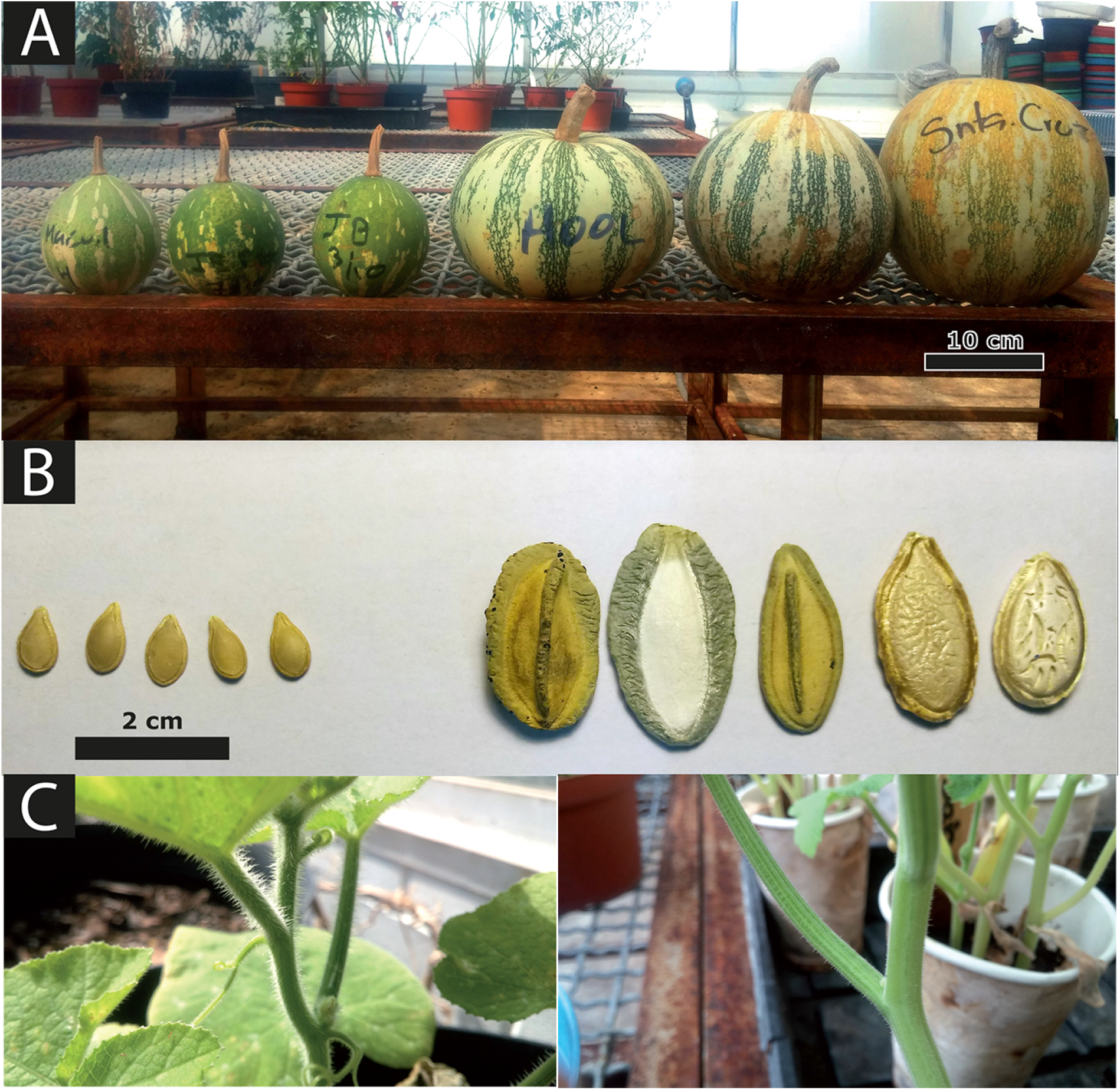
Some morphological differences between *C. argyrosperma* subsp. *sororia* (left) and *C. argyrosperma* subsp. *argyrosperma* (right). Differences in fruit size (A), seed size and shape (B), and in the presence of urticating trichomes (C).

Studies that address the genetic basis of *Cucurbita* domestication are a necessary step towards a deeper understanding of the history of these economically important crops, as well as the development of new breeding strategies and the effective conservation of its genomic resources (Lira et al., 2016; Chomicki et al., 2019). To contribute in this direction, we report here the first genome assembly of the wild relative *sororia*, which complements the existing assembly of *argyrosperma* (Barrera-Redondo et al., 2019). The comparison between the genomes allowed us to find genomic structural variants between both subspecies. We characterized a large sample of *argyrosperma* landraces (117 individuals from 19 locations) and *sororia* accessions (50 individuals from 4 locations) using genome-wide data to investigate their demographic history and propose a domestication scenario. We also performed selection scans throughout the genome of *C. argyrosperma* to detect candidate regions associated with the domestication of this species.

## Results

### Genome assembly of *C. argyrosperma* subsp. *sororia*

We sequenced the genome of a wild individual of subspecies *sororia* using Illumina HiSeq4000 (213x coverage) and PacBio Sequel (75.4x coverage). The genome was assembled in 828 contigs with an N50 contig size of 1.3 Mbp and an L50 of 58 contigs (Table S1). A BUSCO analysis (Simão et al., 2015) against the *embryophyta odb9* database detected 92.8% of complete BUSCOs, 1.2% fragmented BUSCOs and 6.0% missing BUSCOs within the genome assembly, similarly to other *Cucurbita* genome assemblies (Barrera-Redondo et al., 2019; Sun et al., 2017). We predicted 30,694 protein-coding genes within the genome assembly using BRAKER2 (Hoff et al., 2016). Around 35.8% of our *sororia* genome assembly is composed of transposable elements (TEs), slightly higher than the 34.1% of TEs found in a previous *argyrosperma* assembly (Barrera-Redondo et al., 2019).

The genome assembly of *argyrosperma* was previously assembled in 920 scaffolds (Barrera-Redondo et al., 2019), so we aimed to reach chromosome-level assemblies for both the *sororia* and the *argyrosperma* genomes. We anchored 99.97% of the *argyrosperma* genome assembly and 98.8% of the *sororia* genome assembly into 20 pseudomolecules using RaGOO (Alonge et al., 2019), which corresponds to the haploid chromosome number in *Cucurbita* (Whitaker and Bemis, 1975). Both assemblies show high synteny conservation across the genus (Fig. S1) and confirm a previously reported inversion in chromosome four that is shared with *C. moschata* (Sun et al., 2017).

### Structural variants between the wild and domesticated genomes

We compared the genome of *argyrosperma* against the genome of *sororia* (Fig. 2). Some of the centromeres in the genome of *sororia* were larger than in *argyrosperma*, possibly due to a better assembly of the repetitive regions. We found several structural variants (SVs) such as copy-number variants (CNVs), inversions, translocations and unalignable regions between the wild and the domesticated genomes (Table S2). The size of the *sororia* genome assembly is ~254 Mbp, 9.23% larger than the genome assembly of *argyrosperma* (Barrera-Redondo et al., 2019), which could be partially explained by these SVs. The genes found within the CNV losses in *argyrosperma* were enriched in pectinesterases and microtubule-based processes (Table S2). We also found sucrose-6F-phosphate phosphohydrolase (*SPP*) within a CNV loss in *argyrosperma*. The genomes of *argyrosperma* and *sororia* share some common genes within their unalignable regions, such as microtubule-associated proteins and genes related to tryptophan biosynthesis (Table S2), suggesting that those regions contain highly divergent sequences and are not limited to presence/absence variants. However, other unaligned regions contain more genes in *sororia* than in *argyrosperma*, including some proteolytic enzymes and sucrose biosynthetic genes that are absent in the *argyrosperma* genome (Table S2), suggesting that presence/absence variants are also included within the unalignable regions.

**Figure 2.**
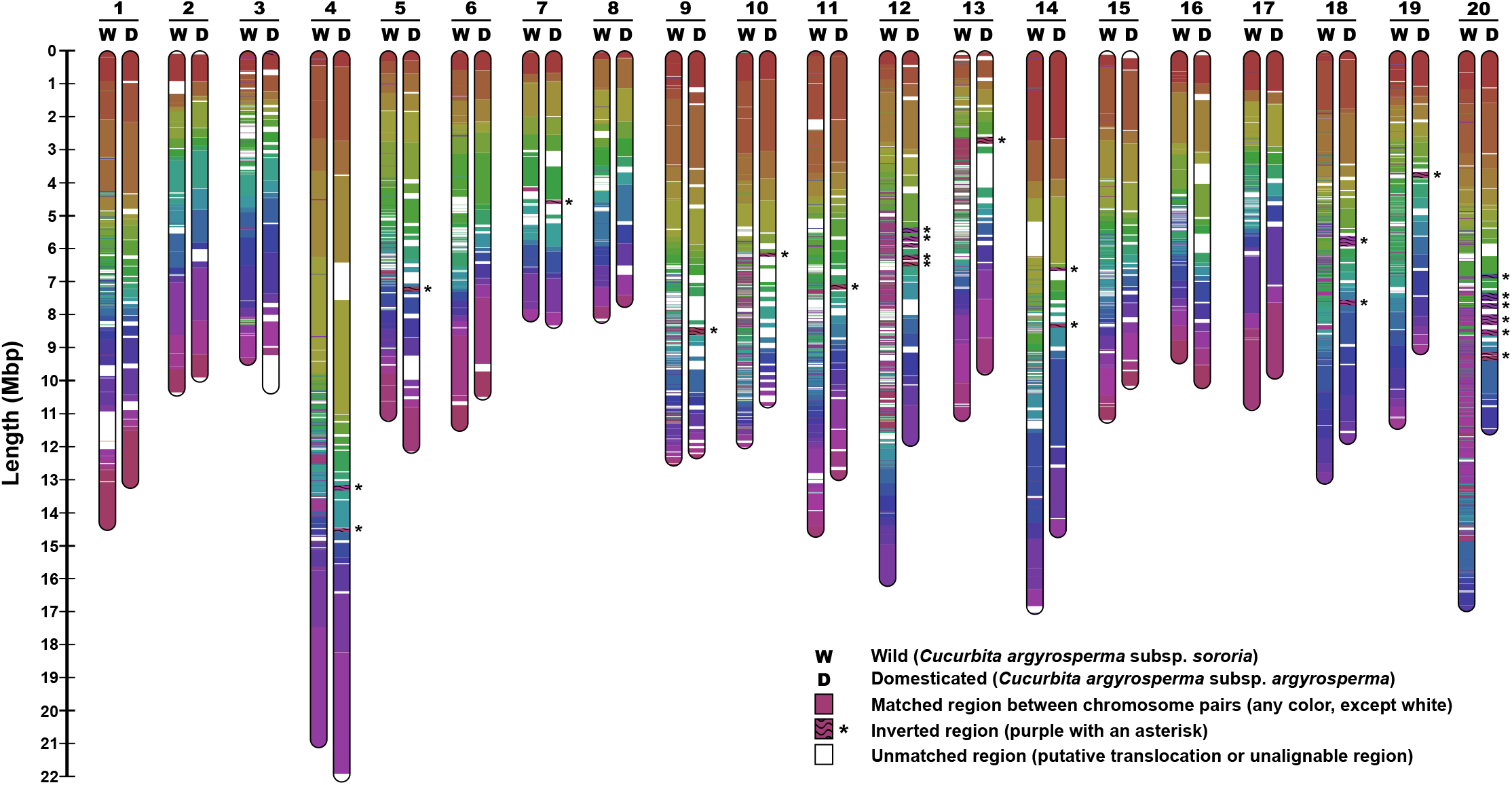
Chromosome map representing the matching regions and putative structural variants between the genome assemblies of *C. argyrosperma* subsp. *sororia* (wild) and *C. argyrosperma* subsp. *argyrosperma* (domesticated). Matching colors represent the aligned homologous regions between both genomes, while white segments represent regions that could not be aligned to the other genome. Inverted regions are highlighted with an asterisk.

### Population data and SNP genotyping

We used samples previously collected throughout Mexico (Sánchez-de la Vega et al., 2018) corresponding to 117 individuals of *argyrosperma*, 50 individuals of *sororia*, 19 feral individuals of *argyrosperma* previously reported to have a semi-wild phenotype and a cultivated genotype based on microsatellite data (Sánchez-de la Vega et al., 2018) and 6 individuals of *C. moschata* (the domesticated sister species of *C. argyrosperma*), that were used as outgroups (Table S3). The samples were sequenced using the tunable Genotype by Sequencing (tGBS) method (Ott et al., 2017) to obtain genome-wide genetic information of the *C. argyrosperma* populations. The reads were quality-filtered and mapped against the chromosome-level genome assemblies of *argyrosperma* and *sororia* to predict single nucleotide polymorphisms (SNPs) and assess possible reference biases in the SNP prediction. Using the reference genome of *argyrosperma*, we obtained an initial dataset consisting of 12,813 biallelic SNPs with a mean read depth of 50 reads per SNP and a minor allele frequency (MAF) of at least 1% (13k dataset, Dataset S1). We also mapped the whole-genome Illumina reads of *argyrosperma* and *sororia*, as well as the whole-genome sequencing of a *C. moschata* individual and a *C. okeechobeensis* subsp. *martinezii* individual (a closely related wild *Cucurbita* species), against the reference genome of *argyrosperma* to obtain a dataset of 11,498,421 oriented biallelic variants (SNPs and indels) across the genome that was used to assess introgression and incomplete lineage sorting (Dataset S2).

### Demographic history of *C. argyrosperma* during its domestication

We eliminated the SNPs that deviated from Hardy-Weinberg equilibrium (exact test with *p* < 0.01) and pruned nearby SNPs under linkage disequilibrium (LD with an r^2^ > 0.25 in 100 kbp sliding windows) from the 13k dataset to retrieve a set of 2,861 independent SNPs that could be used for demographic analyses.

We found similar genetic variation in *sororia* and *argyrosperma*, regardless of the reference genome used (average nucleotide diversity *π* range 0.095 - 0.098 for both taxa, see Table S4). At a population scale, the wild population in Jalisco had the highest genetic diversity within *sororia*, while the highest diversity in *argyrosperma* was found in the Pacific Coast of Mexico (Table S5). The domesticated and wild populations of *C. argyrosperma* displayed low genetic differentiation (*F*_ST_ = 0.0646; 95% confidence interval from 0.0565 to 0.0751), while feral populations were more closely related to *argyrosperma* (*F*_ST_ = 0.0479) than with *sororia* (*F*_ST_ = 0.1006).

We used SNPhylo (Lee et al., 2014) and ADMIXTURE (Alexander et al., 2009) to evaluate the genealogical relationships and genetic structure among the wild and domesticated populations of *C. argyrosperma*. We confirmed the genetic differentiation between *sororia* and *argyrosperma*, as detected by the *F*_ST_ analyses. Our Maximum Likelihood (ML) tree groups all the *argyrosperma* populations in a single monophyletic clade (Fig. 3A). We found additional genetic differentiation between the *sororia* populations in Southern Mexico (populations 1-3) and the *sororia* populations in Jalisco (population 4), in both the ADMIXTURE assignations (Fig. 3B) and their positions in the ML tree (Fig. 3A). The *sororia* populations of Jalisco are genetically closer to *argyrosperma*, as shown by their paraphyletic position in the ML tree (Fig. 3A). Consistent with a domestication in the lowlands of Western Mexico, the *argyrosperma* populations of Guerrero and Jalisco represent the basal branches of the *argyrosperma* clade (Fig. 3A), all showing instances of genetic similarity to the *sororia* populations in Jalisco in the four genetic groups (K) of ADMIXTURE (Fig. 3B). The *argyrosperma* populations in Western Mexico (populations 5-17) are genetically differentiated from the Eastern populations (populations 18-26), with a possible recent anthropogenic dispersion event of Eastern populations into Onavas, Sonora (population 19; Fig. 3B-C). These four genetic groups are also retrieved in a principal component analysis (PCA; Fig. S2). The ADMIXTURE results (Fig. 3B) uncovered admixture events between some *sororia* and *argyrosperma* populations. This pattern is evident in the populations of Oaxaca and Sinaloa (populations 14 and 18; Fig. 3C). The feral populations are consistently grouped alongside their sympatric domesticated populations within *argyrosperma* (Fig. 3A-B), indicating that these populations diverged recently from nearby domesticated populations.

**Figure 3.**
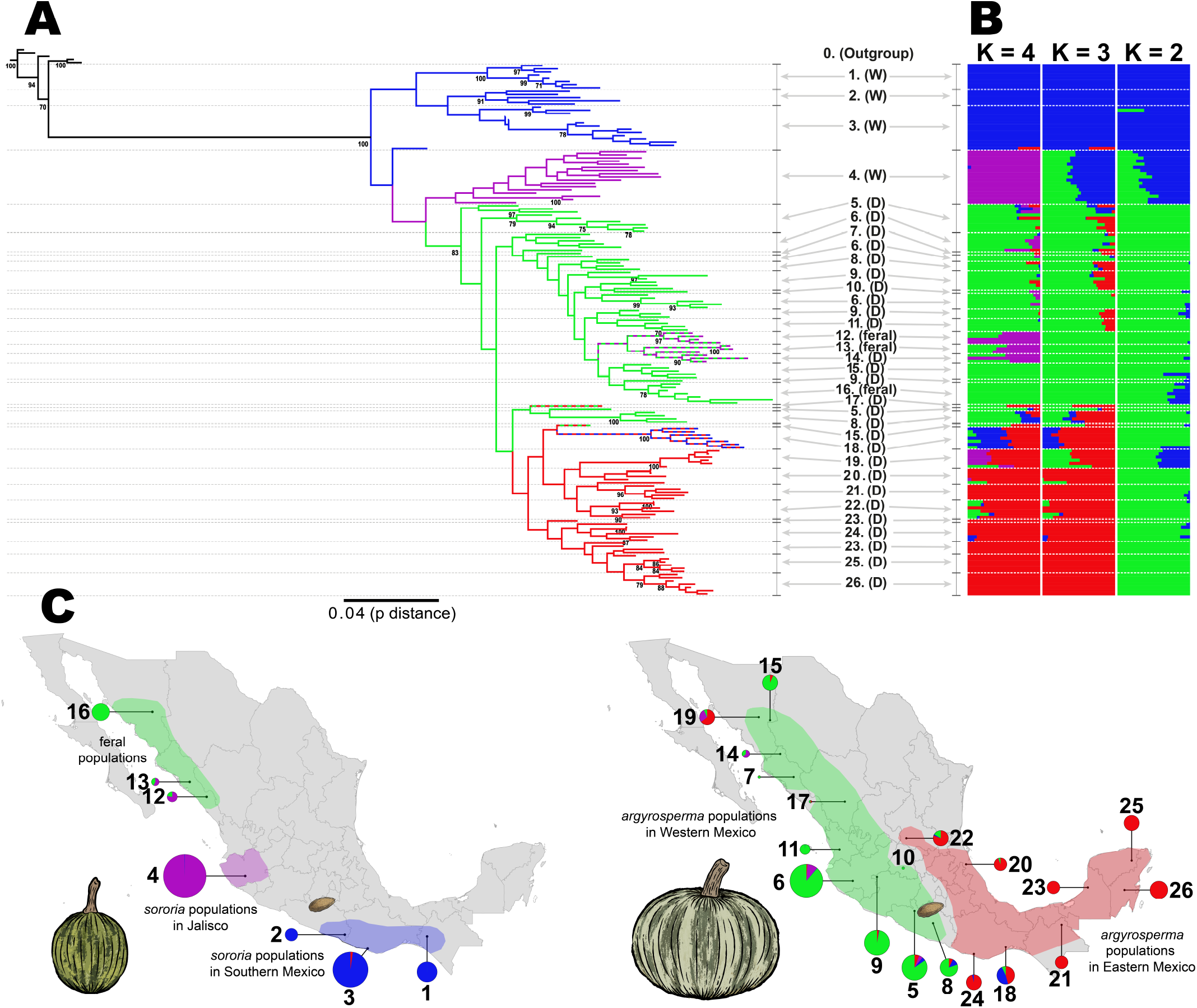
Genetic structure and phylogenetic relationships between the wild and domesticated populations of *Cucurbita argyrosperma* based on 2,861 SNPs. (A) Maximum Likelihood tree with 100 bootstraps (only bootstrap values > 70 shown) (B) ADMIXTURE analysis using K values ranging from 2 to 4. (C) Geographic distribution of the wild (down left) and domesticated (down right) populations, with pie chart colors representing the ADMIXTURE assignation of the individuals in 4K ancestral populations (size of pie charts proportional to sample size). The seed in the maps represent the earliest archaeological record of *argyrosperma* from Xihuatoxtla, Guerrero (dated 8,700 years BP) (Piperno et al., 2009).

We used Fastsimcoal 2 (Excoffier and Foll, 2011) to test whether *argyrosperma* was domesticated from a *sororia* population in Southern Mexico or from a *sororia* population in Jalisco. Given that gene flow has been previously observed between *argyrosperma* and *sororia* (Montes-Hernandez and Eguiarte, 2002), we compared three different gene flow models (continuous gene flow, secondary contact or no gene flow) for each scenario (Fig. 4A). A comparison between models using the Akaike Information Criterion indicates that the Jalisco domestication model with secondary contact (*i.e*., extant gene flow after initial genetic isolation between subspecies) is the most likely of the domestication scenarios assayed.

**Figure 4.**
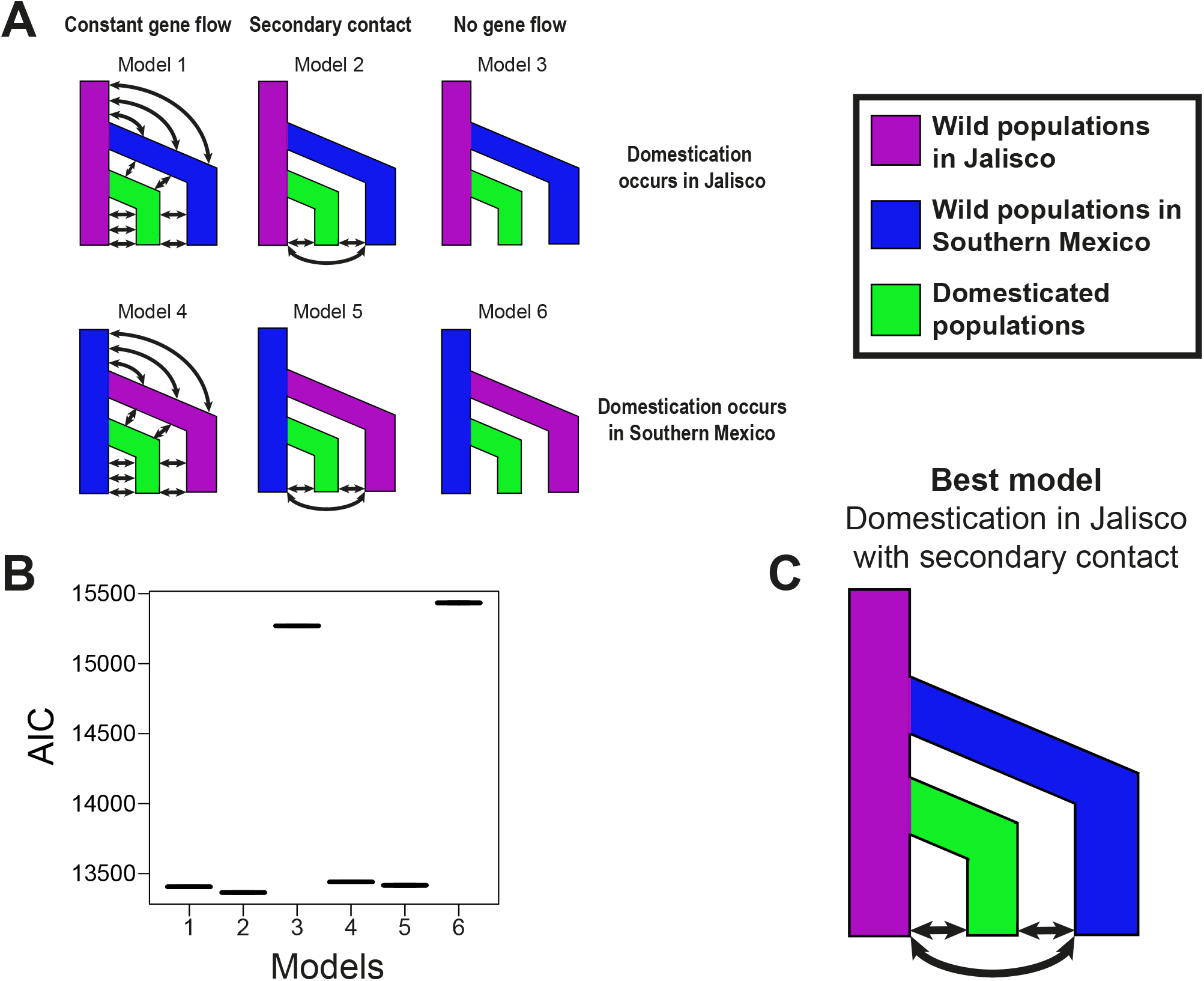
Coalescent simulations and most likely domestication scenario of *Cucurbita argyrosperma*. (A) The six models assessed against the unfolded multidimensional Site Frequency Spectrum of our data. (B) Comparison of the Akaike Information Criterion (AIC) of all the models. (C) The domestication model that best fits the data.

### Domestication sweeps in *C. argyrosperma*

In order to perform the tests to detect selective sweeps associated with the domestication of *C. argyrosperma*, we removed from the 13k dataset the *C. moschata* individuals as well as the feral individuals of *C. argyrosperma*. We used the 1% MAF threshold for this subset, obtaining a 10,617 SNP dataset suitable to detect selective sweeps, with a marker density of 44 SNPs per Mb. LD was limited within the dataset, with a mean pairwise r^2^ of 0.1 (Fig. S3). We performed two *F*_ST_-based tests as implemented in BayeScEnv (de Villemereuil and Gaggiotti, 2015) and PCAdapt (Luu et al., 2017) to detect selective sweeps between the domesticated and the wild populations of *C. argyrosperma* (Fig. 5A). BayeScEnv is an *F*_ST_-based method that tests correlation with other variables, in our case the wild or domesticated nature of each population (coded as 0 and 1, respectively). PCAdapt does not require an *a priori* grouping of individuals into wild/domesticated, as we used the two principal components of a PCA to control for the underlying genetic structure between subspecies (Fig. S2).

**Figure 5.**
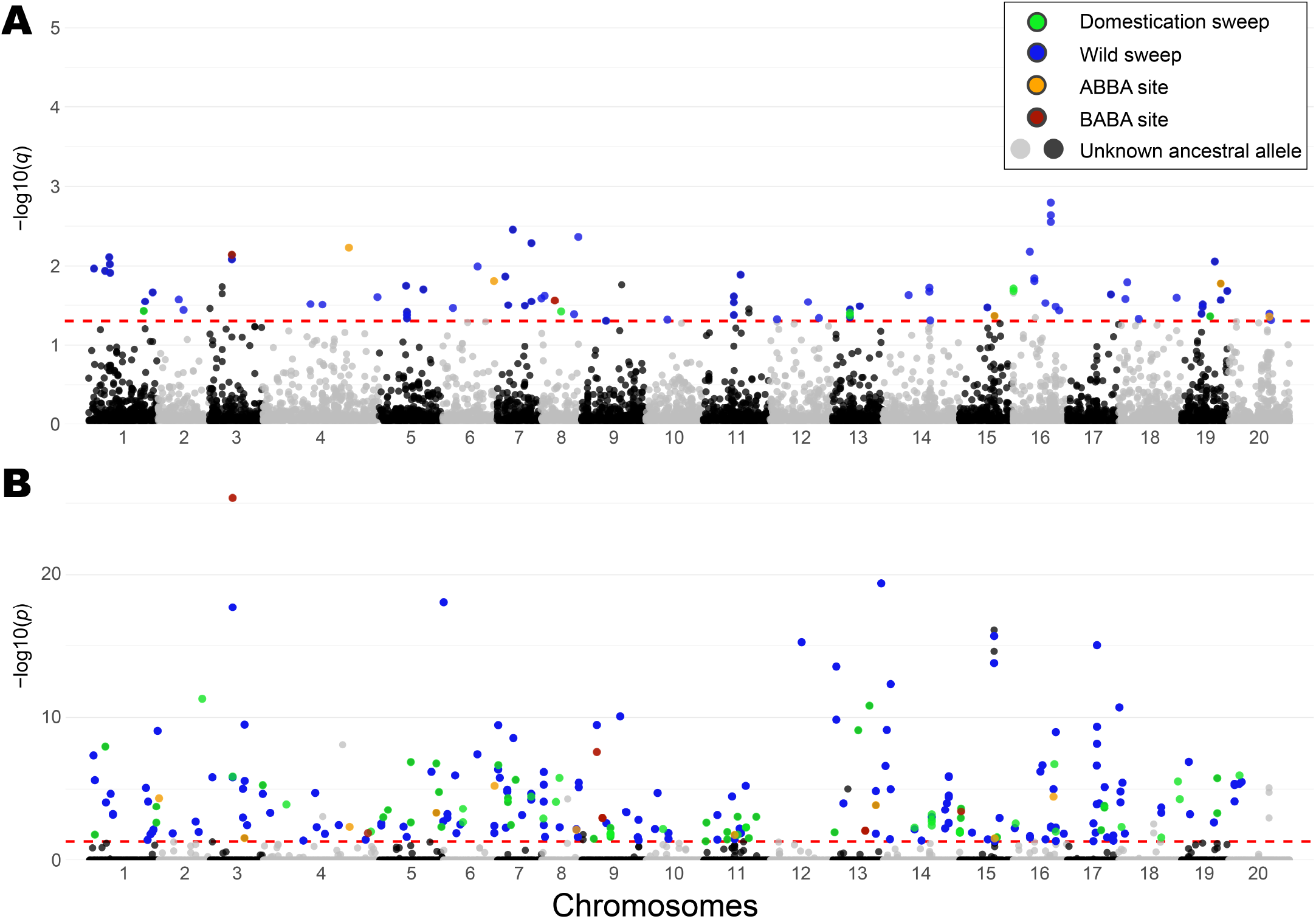
Putative footprints of selection associated with the domestication of *Cucurbita argyrosperma*. Manhattan plots representing the (A) BayeScEnv and (B) PCAdapt tests in each chromosome of the genome. The red line indicates the cutoff value (*q*-value or Bonferroni-corrected *p*-value < 0.05) to determine a candidate locus. The colors of the outlier loci indicate whether the putative selective signal corresponds to *argyrosperma* (green), *sororia* (blue), an ABBA site (orange) a BABA site (red) or an unknown selective direction (grey or black dots above threshold).

We discovered 338 outlier SNPs with either BayeScEnv (91 outliers, Fig. 5A) or PCAdapt (266 outliers, Fig. 5B). We used *C. moschata* and *C. okeechobeensis* subsp. *martinezii* as outgroups to determine the direction of the putative selective pressures for each SNP outlier, as well as determining possible introgression or incomplete lineage sorting between *sororia*, *argyrosperma* and *C. moschata*. We found that 81 of the outliers corresponded to selective signals in *argyrosperma*, while 215 outliers corresponded to selective signals in *sororia* (Fig. 5). We could not determine the direction of selection for 22 SNP outliers, and 20 outliers showed signals of introgression or incomplete lineage sorting. Only 19 SNPs were shared as outliers by both tests, from which 15 corresponded to selective signals in *sororia* and 4 showed putative signals of introgression.

We identified several instances of either genetic introgression or incomplete lineage sorting between *C. moschata* and both *argyrosperma* (ABBA sites) or *sororia* (BABA sites) (Fig. S4). We performed a genome-wide *D*-statistic analysis and found significantly more instances of shared derived variants between *argyrosperma* and *C. moschata* than between *sororia* and *C. moschata* (block-jackknife *p*-value = 0.0014), obtaining an overall admixture fraction *f*_G_ of 0.01 (Table S6). From the 20 outliers with putative signals of introgression, 12 correspond to ABBA sites and 8 correspond to BABA sites (Fig. 5).

We found 125 protein-coding genes and 7 long-noncoding RNAs including outlier SNPs within their structure (*i.e.*, introns, exons, UTRs), which were assigned according to the observed direction of the putative selective signals (Table S7). Among the genes under putative selection in *sororia* were a homolog of Auxin response factor 1 (*ARF1*), ABC transporter C family member 2 and six serine/threonine-protein kinases (Table S7). Among the genes under putative selection in *argyrosperma*, we found transcription factor *MYB44*, Glycerophosphodiester phosphodiesterase *GDPDL4*, auxin-responsive protein *IAA27*, *WRKY* transcription factor 2 (*WRKY2*), ABC transporter E family member 2, Filament-like plant protein 4 (*FPP4*), the ATP-dependent zinc metalloprotease *FTSH11*, *DMR6*-like oxygenase 1 (*DLO1*) and serine/threonine-protein kinase *CES101* (Table S7). We also found genes under putative selection overlapping ABBA and BABA sites (Table S7). We found a homolog of flowering locus K (*FLK*) under putative selection in *argyrosperma* and as an ABBA site. We also found a transport inhibitor response 1 (*TIR1*) homolog under putative selection in both *sororia* and as a BABA site.

After performing a Gene Ontology enrichment analysis (Alexa et al., 2006), we found seven significantly enriched biological and biochemical processes in the 125 candidate genes, including abscisic acid (ABA) biosynthesis and brassinosteroid-mediated signaling (Table S8). The zeaxanthin epoxidase activity was also significantly enriched in the 125 candidate genes (Table S8).

We identified 79 candidate SNPs that resided within 59 of the SVs found between the genome assemblies of *sororia* and *argyrosperma* (Table S9). Of these SVs, 10 correspond to putative selective signals in *argyrosperma*, suggesting they could be associated with its domestication. One CNV gain contains four long noncoding RNAs and additional copies of the mitochondrial genes *COX3*, *ND2* and *YMF19*. Another CNV contains the gene *MKP1*, which was found under putative selection in both *argyrosperma* as a copy loss and in *sororia* as a copy gain. A third SV consists of a translocation containing an uncharacterized protein. Interestingly, the rest of these domestication SVs only contain long noncoding RNAs (two CNVs and one unaligned region containing 9 long noncoding RNAs), tRNAs (one CNV with 5 tRNAs), or intergenic DNA (two translocations and one CNV). We also found three SVs corresponding to ABBA sites that represent putative introgression signals between *argyrosperma* and *C. moschata*. One SV is a CNV that contains an additional copy of a Ras-related protein *Rab7* near its candidate SNP (which was identified as an outlier by both PCAdapt and BayeScEnv), another is a translocation that contains the non-specific lipid-transfer protein *AKCS9* near its candidate SNP, and the third is an unalignable region containing tRNAs.

## Discussion

The genome assembly of *sororia* represents the first sequenced genome of a wild *Cucurbita*, which allowed us to detect structural and functional differences with the *argyrosperma* genome. The genome assembly of *argyrosperma* was smaller than the *sororia* assembly, which is possibly caused by the loss of structural variants during its domestication, as has been reported in pan-genome studies (Khan et al., 2020). Many of these unaligned regions contain entire genes in *sororia*, making this wild taxon a reservoir of potentially adaptive presence/absence variants. However, the extant genetic diversity of *argyrosperma* is similar to that of *sororia*, which suggests that the effects of the domestication bottleneck were alleviated by the current gene flow between both subspecies as suggested by our coalescent simulations and by the results of previous studies (Montes-Hernandez and Eguiarte, 2002). This gene flow may be related to the sympatric distribution of the wild and domesticated populations of *C. argyrosperma* throughout the Pacific Coast of Mexico (Sánchez-de la Vega et al., 2018), where their coevolved pollinator bees *Peponapsis* spp. and *Xenoglossa* spp. are found (Wilson, 1990). Traditional agricultural practices are another fundamental force that maintains the diversity of crop species (Jarvis et al., 2008). Since *argyrosperma* is a traditional crop cultivated for both self-supply and local markets where it has a specialized gastronomic niche (Lira et al., 2016), the genetic diversity in *argyrosperma* is also maintained by the conservation of local landrace varieties at local scales (Montes-Hernández et al., 2005; Barrera-Redondo et al., 2020).

Our demographic analyses suggest that the extant populations from Jalisco are the closest modern relatives of the initial population of *sororia* from which *argyrosperma* originated. The genetic relatedness between the *sororia* populations from Western Mexico and *argyrosperma* was also observed with mitochondrial markers (Sanjur et al., 2002). The domestication of *C. argyrosperma* likely started around 8,700 years ago, as suggested by the earliest, albeit taxonomically ambiguous, archaeological record of *argyrosperma* (Piperno et al., 2009; Ranere et al., 2009). Since crop domestication in Mesoamerica is linked to migration patterns and cultural development of early human populations in America (Zizumbo-Villarreal and Colunga-GarcíaMarín, 2010; Zizumbo-Villarreal et al., 2012), we expected *argyrosperma* to share historical demographical patterns with human history. Our data shows that the *argyrosperma* populations found in Guerrero and Jalisco are the most closely related to the *sororia* populations from Jalisco. This means that even if the closest wild relatives of *argyrosperma* are currently found in Jalisco, the domestication process may have occurred throughout the lowlands of Jalisco and the Balsas basin (Piperno et al., 2009). Ancient human migration events have been proposed to occur along the river basins in Southwestern Mexico, which may explain the genetic cohesiveness among the *argyrosperma* populations of that area that represent the first fully domesticated lineage of the species (Zizumbo-Villarreal and Colunga-GarcíaMarín, 2010). Previous studies based on 8,700 years old phytoliths found in Xihuatoxtla, Guerrero, suggest the co-occurrence of *Zea mays* and *C. argyrosperma* in the Balsas region within this time period (Piperno et al., 2009; Ranere et al., 2009). Overall, the patterns of genetic structure for *C. argyrosperma* are coherent with the archaeological evidence of early human migration throughout Mesoamerica (Stinnesbeck et al., 2017; Piperno, 2011) (Fig. 6).

**Figure 6.**
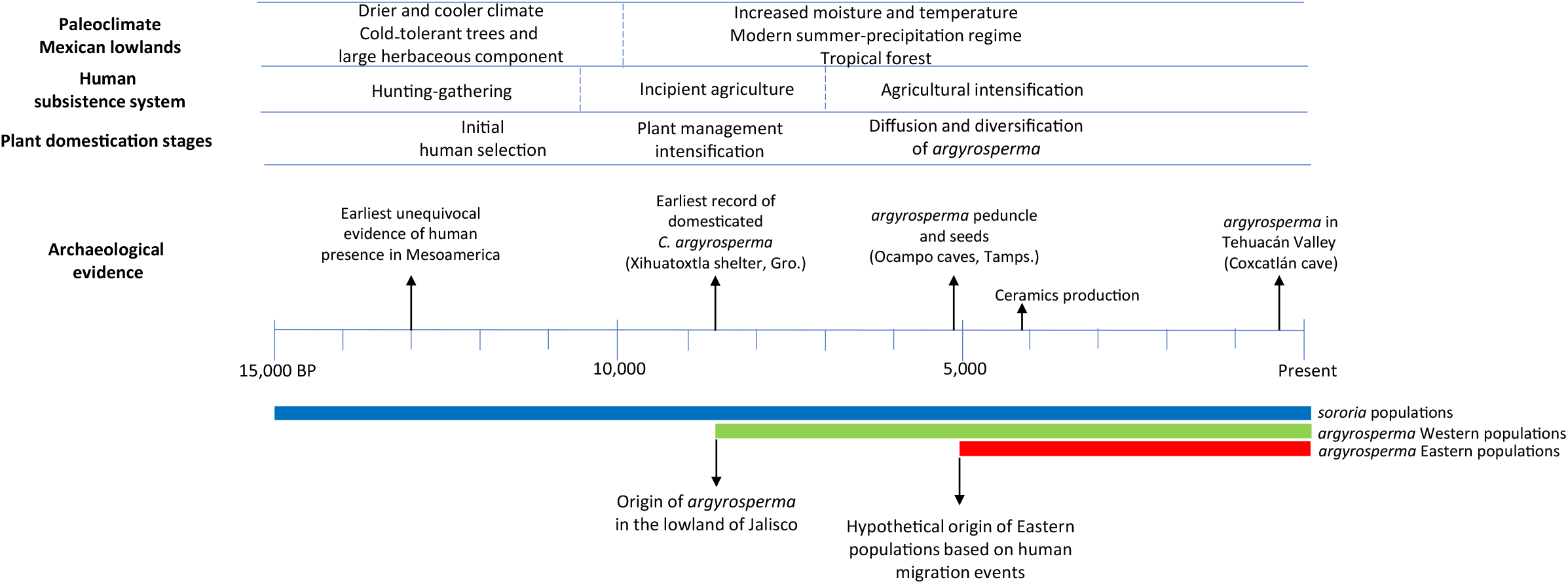
Model of *Cucurbita argyrosperma* domestication based on archaeological and genetic data (Smith 1997; Sanjur et al., 2002; Ranere et al., 2009; Zizumbo-Villarreal and Colunga-GarcíaMarín, 2010; Stinnesbeck et al., 2017). The colored lines represent the presence of each *C. argyrosperma* population in Mesoamerica (blue = *sororia* populations, green = *argyrosperma* populations in Western Mexico, red = *argyrosperma* populations in Eastern Mexico).

We found several signals of putative selection between *argyrosperma* and *sororia*, even though tGBS sequencing has a limited capacity to detect selective sweeps across the genome (Lowry et al., 2017). The SNP density for our selection tests was of 44 SNPs per Mb, which is one order of magnitude denser than other studies using reduced-representation genome sequencing to detect selective sweeps (Lowry et al., 2017). This is a consequence of the relatively small genome size of *C. argyrosperma* (Barrera-Redondo et al., 2019). Nonetheless, the LD in *C. argyrosperma* decays at a shorter length than our SNP density, so our genome scans should be interpreted as a partial representation of the selective sweeps associated with the domestication process (Lowry et al., 2017).

Most of the putative selective signals were attributed to *sororia*, probably because wild taxa are subject to many natural selective pressures. Many of the SNPs that were retrieved as outliers were found on genes involved in biotic and abiotic plant defense responses. For example, *PBL10* and *PBL23* have been suggested to be involved in plant defense pathways due to their similarity to other serine/threonine-protein kinases (Zhang et al., 2010).

However, we also found some genes under putative selection in *argyrosperma* that could be attributed to defense mechanisms such as *MYB44* and *DLO1*, involved in several defense responses (Shim and Choi, 2013; Zeilmaker et al., 2015); heat response with *FTSH11* (Chen et al., 2006); and trichome morphogenesis and differentiation with *FPP4* and *GDPDL4* (Hayashi et al., 2008; Chen et al., 2016), which is a morphological characteristic that differentiates *argyrosperma* from *sororia* (Fig. 1). This is concordant with previous studies showing that selective pressures during domestication actively purge these defense mechanisms, as the products of these responses are usually unpleasant or harmful to humans when the plant is consumed (Moreira et al., 2018). This is particularly important for breeding programs, since wild *Cucurbita* such as *sororia* harbor loci associated to disease resistance that their domesticated counterparts have lost (Paris, 2016). The selective pressures found in *MKP1* suggest a disruptive selection regime between *sororia* and *argyrosperma*. Since *MKP1* modulates defense responses (Ulm et al., 2002), it is possible that both subspecies adapted to differential environmental pressures as domestication took place.

Our Gene Ontology enrichment analysis found several genes enriched in the ABA biosynthesis and brassinosteroid-mediated signaling. This suggests that the alteration of growth hormones may play an important role in *C. argyrosperma* domestication. ABA is involved in a myriad of functions, such as the regulation of plant growth, plant development, seed dormancy and response to biotic/abiotic stress (Chen et al., 2020). In this sense, the lack of dormancy in seeds and gigantism are both common domestication changes that are present in domesticated cucurbits that may be caused by changes in the regulation of ABA and brassinosteroids (Martínez et al., 2017; Chomicki et al., 2019). Particularly, phytohormone regulation may be involved in *Cucurbita* fruit size alongside microtubule-related genes (Chomicki et al., 2019), such as the microtubule-associated proteins we found in our structural variant analysis. We found *WRKY2* among our candidate genes, which is involved in seed germination and post-germination development in *A. thaliana* (Jiang and Yu, 2009) and may explain the lack of seed dormancy in *argyrosperma*. Likewise, we found *IAA27* under selection in *argyrosperma*, which is involved in plant growth and development (Liscum and Reed, 2002).

Among specific candidate genes, we found two ABC transporters under selection in both *argyrosperma* and *sororia*. Some ABC transporters are involved in the transmembrane transport of ABA-GE, an ABA conjugate that is usually attributed to the plant response against water stress (Burla et al., 2013). However, previous studies in *Hordeum vulgare* suggest that the transport of ABA-GE may play a role in seed development alongside *de novo* ABA synthesis within the developing seed (Seiler et al., 2011), suggesting a role of ABC transporters in the seed development of *C. argyrosperma*. Previous studies have also identified an association between variants in ABC transporter proteins and seed size in *Cucurbita maxima* (Wang et al., 2019) and *Linum usitatissimum* (Guo et al., 2019), further suggesting that ABA may be deregulated via selective pressures on ABC transporters to enhance seed size in *C. argyrosperma* during its domestication. Variants in a serine/threonine-protein kinase, as the ones we found in our selective scans, have also been associated with seed size in *C. maxima* (Wang et al., 2019). We found *SPP* as a CNV loss in *argyrosperma*, whose ortholog in *C. maxima* has been predicted as a major QTL for seed length (Wang et al., 2019). We also detected a significant enrichment of zeaxanthin epoxidase activity. Zeaxanthin epoxidase is linked to the degradation of carotenoids in plant seeds (Gonzalez-Jorge et al., 2016), which in turn are important precursors of ABA biosynthesis, regulating processes such as germination and maturation (Frey et al., 2006).

We found shared derived variants between *C. moschata* and *argyrosperma* under putative selection. We located *FLK*, which is involved in flowering time regulation (Mockler et al., 2004), under selection in *argyrosperma* and as an ABBA site (*i.e.*, as a putatively introgressed allele between both domesticated taxa). We also found *Rab7* and *AKCS9* within ABBA-related SVs, which are involved in processes such as plant growth and seed development (Cui et al., 2014; Liu et al., 2015). This suggests that, given the close relationship between *C. argyrosperma* and *C. moschata*, both species may share domestication loci involved in common domestication traits such as flowering time, plant growth and seed growth. However, we also found *TIR1* under selection in *sororia* and as a BABA site, which is an auxin receptor involved in ethylene signaling and antibacterial resistance in roots (Navarro et al., 2006). These ABBA and BABA sites under selection may be shared with *C. moschata* either due to incomplete lineage sorting or by adaptive introgression with the wild and domesticated populations of *C. argyrosperma*. The incorporation of domesticated loci between *argyrosperma* and *C. moschata* through introgression may have been an effective way for Mesoamerican cultures to domesticate multiple *Cucurbita* taxa. This hypothesis is supported by the significant amount of ABBA sites we found between the genomes of *argyrosperma* and *C. moschata*. However, this hypothesis needs to be further addressed using population-level data of *C. moschata* and other domesticated *Cucurbita* species.

## Methods

### Genome assembly and annotation of *Cucurbita argyrosperma* subsp. *sororia*

We sequenced and assembled *de novo* the genome of a *sororia* individual collected in Puerto Escondido (Oaxaca, Mexico). Its DNA was extracted from leaf tissue and sequenced using PacBio Sequel at the University of Washington PacBio Sequencing Services and using Illumina HiSeq4000 at the Vincent J. Coates Genomics Sequencing Laboratory in UC Berkeley (NIH S10 Instrumentation Grants S10RR029668 and S10RR027303). We filtered the Illumina sequences using the qualityControl.py script (https://github.com/Czh3/NGSTools) to retain the reads with a PHRED quality ≥ 30 in 85% of the sequence and an average PHRED quality ≥ 25. The Illumina adapters were removed using SeqPrep (https://github.com/jstjohn/SeqPrep) and the paired reads that showed overlap were merged. The chloroplast genome was assembled using NOVOplasty (Dierckxsens et al., 2017) and the organellar reads were filtered using Hisat2 (Kim et al., 2019) against the chloroplast genome of *argyrosperma* (Barrera-Redondo et al., 2019) and the mitochondrial genome of *C. pepo* (Alverson et al., 2010). We assembled the nuclear genome into small contigs using the Illumina reads and the Platanus assembler (Kajitani et al., 2014). The Platanus contigs were assembled into larger contigs using the PacBio Sequel reads and DBG2OLC (Ye et al., 2016). We performed two iterations of minimap2 and racon (Li, 2018) to obtain a consensus genome assembly by mapping the PacBio reads and the Platanus contigs against the DBG2OLC backbone. We performed three additional polishing steps using PILON (Walker et al., 2014) by mapping the Illumina reads against the consensus genome assembly with BWA *mem* (Li and Durbin, 2010).

The genome annotation processes were performed using the GenSAS v6.0 online platform (Humann et al., 2019). The transposable elements within the genome were predicted and masked using RepeatModeler (http://www.repeatmasker.org/RepeatModeler/). We downloaded five RNA-seq libraries of *C. argyrosperma* available on the Sequence Read Archive (accessions SRR7685400, SRR7685404 - SRR7685407) to use them as RNA-seq evidence for the gene prediction. We performed the same quality filters described above for the RNA-seq data and aligned the high-quality reads against the masked genome of *C. argyrosperma* subsp. *sororia* using STAR v2.7 (Dobin et al., 2013). We used filterBAM from the Augustus repository (Stanke et al., 2006) to filter low-quality alignments and used the remaining alignments as RNA-seq evidence to predict the gene models using BRAKER2 (Hoff et al., 2016). The gene predictions were functionally annotated using InterProScan (Jones et al., 2014) and by aligning the gene models against the SwissProt database (Schneider et al., 2009) using BLASTp (Camacho et al., 2009) with an *e*-value < 1e^−6^.

### Anchoring the reference genomes into pseudomolecules

We aimed to improve the genome assembly of *argyrosperma*, which was previously assembled in 920 scaffolds (Barrera-Redondo et al., 2019), and reach chromosome-level assemblies for both genomes. Thus, we generated Pacbio corrected reads from the published PacBio RSII reads of *argyrosperma* (NCBI SRA accession SRR7685401) and the PacBio Sequel reads of *sororia* (sequenced for this study at the University of Washington PacBio Sequencing Services) using CANU (Koren et al., 2017). We anchored the genome assemblies of *argyrosperma* (Barrera-Redondo et al., 2019) and *sororia* into pseudomolecules using RaGOO (Alonge et al., 2019) alongside the PacBio corrected reads of each taxon to detect and correct misassemblies, using a gap size of 2600 bp for chromosome padding (corresponding to the average gap length of the *argyrosperma* genome assembly) and using the genome assembly of *C. moschata* (Sun et al., 2017) as reference. The chromosome numbers in both assemblies were assigned in correspondence to the genome assembly of *C. moschata* (Sun et al., 2017).

### Structural variant analysis

We evaluated the synteny between *Cucurbita* genomes using Synmap2 (Haug-Baltzell et al., 2017). We analyzed the genome rearrangements and structural variants between *sororia* and *argyrosperma* using Smash (Pratas et al., 2015) with a minimum block size of 100,000 bp, a threshold of 1.9 and a context of 28; and using SyRI (Goel et al., 2019) alongside nucmer (Kurtz et al., 2004) with a minimum cluster length of 500 bp, an alignment extension length of 500 bp, a minimum match length of 100 bp and a minimum alignment identity of 90%. The gene content associated with each type of structural variant was considered either as the overlap between genes and variants (inversions and translocations) or as the genes contained entirely within the structural variants (copy-number variants and unaligned regions). We performed a Gene Ontology enrichment analysis using topGO and the *weight01* algorithm (Alexa et al., 2006) to find enriched biological functions associated to each type of structural variant. We determined the significantly enriched biological functions by performing Fisher’s exact test (*p*-value < 0.05).

### Data filtering and SNP genotyping

We used previously collected seeds from 19 populations of *argyrosperma* landraces, four populations of *sororia*, and three feral populations (Sánchez-de la Vega et al., 2018), covering most of the reported distribution of this species throughout Mexico (Castellanos-Morales et al., 2018) (Table S3). The seeds were germinated in a greenhouse and total DNA was extracted from fresh leaves using a DNeasy Plant MiniKit (Qiagen) of 192 individuals across the collected populations (Table S3), including five individuals of *C. moschata* to be used as outgroup. All 192 individuals were sequenced by Data2Bio LLC using the tunable Genotyping by Sequencing (tGBS) method (Ott et al., 2017) with an Ion Proton instrument and two restriction enzymes (Sau3AI/BfuCI and NspI). The wild and domesticated populations were randomly assigned to the plate wells before library preparation to avoid sequencing biases.

The raw reads of the tGBS sequencing were trimmed using LUCY2 (Li and Chou, 2004), removing bases with PHRED quality scores < 15 using overlapping sliding windows of 10bp. Trimmed reads shorter than 30 bp were discarded. The trimmed reads were mapped against the chromosome-level genome assembly of *argyrosperma* using segemehl (Hoffmann et al., 2009), since empirical studies suggest this read-mapping software outperforms others for Ion Torrent reads (Caboche et al., 2014). We only retained the reads that mapped uniquely to one site of the reference genome for subsequent analyses.

We used BCFtools (Li et al., 2009) for an initial variant calling step, retaining variants with at least 6 mapped reads per individual per site where the reads had a minimum PHRED quality score of 20 in the called base and a minimum mapping quality score of 20 (Li et al., 2008). We used plink (Purcell et al., 2007) to perform additional filters, such as retaining only biallelic SNPs, retaining SNPs with no more than 50% of missing data, individuals with no more than 50% of missing data and sites with a minor allele frequency (MAF) of at least 1% (13k dataset). After eliminating individuals with missing data, only 109 individuals of *argyrosperma*, 44 individuals of *sororia*, 14 feral individuals and 5 individuals of *C. moschata* remained for the subsequent analyses. We repeated the SNP prediction using the reference genome of *sororia* to evaluate potential reference biases. We found a similar number of SNPs (10,990) and comparable estimates of genetic diversity (see Table S4), suggesting that reference bias does not have a meaningful impact on our results. Thus, we employed the domesticated genome as the reference for the rest of the population analyses.

In order to obtain an adequate SNP dataset to infer the demographic history of *C. argyrosperma*, we performed additional filters to the 13k dataset with plink (Purcell et al., 2007), including the elimination of all the SNPs that diverged significantly (*p* < 0.01) from the Hardy-Weinberg equilibrium exact test (Wigginton et al., 2005), and the elimination of adjacent SNPs with a squared correlation coefficient (r^2^) larger than 0.25 within 100 kbp sliding windows with a step size of 100 bp.

We also generated a SNP dataset to detect selective sweeps associated with the domestication of *C. argyrosperma* by eliminating all the feral individuals of *C. argyrosperma*, which could not be assigned to either a wild or a domesticated population, as well as the five individuals of *C. moschata*. We also eliminated the SNP sites with more than 50% missing data and performed a MAF filter of 1% after reducing the number of individuals in the 13k dataset. The SNP density was calculated with VCFtools (Danecek et al., 2011) and the LD decay was calculated using plink (Purcell et al., 2007) with a minimum r^2^ threshold of 0.001.

We also sequenced the genome of a *C. moschata* individual from Chiapas (Mexico) and the genome of a *C. okeechobeensis* subsp. *martinezii* individual from Coatepec (Veracruz, Mexico) using the Illumina HiSeq4000 platform in UC Berkeley. We downloaded the genome sequences of *argyrosperma* (Barrera-Redondo et al., 2019) from the Sequence Read Archive (accessions SRR7685402 and SRR7685403). The Illumina whole-genome sequences were filtered using the same quality parameters as the ones used in the genome assembly of *sororia* (see above) and were aligned against the chromosome-level assembly of *argyrosperma* using BWA *mem* (Li and Durbin, 2010). We only retained the reads that mapped uniquely to one site of the reference genome and retained only the biallelic sites with a sequencing depth >=10 reads per genome.

### Population structure

We used diveRsity (Keenan et al., 2013) to calculate the pairwise *F*_ST_ statistics, using 100 bootstraps to calculate the 95% confidence intervals. We estimated the genetic variation in the wild, domesticated and feral populations with STACKS (Catchen et al., 2013). Using ADMIXTURE (Alexander et al., 2009), we evaluated the genetic structure among the *sororia* and *argyrosperma* populations, evaluating their individual assignment into one (CV error = 0.26205), two (CV error = 0.25587), three (CV error = 0.25806) and four (CV error = 0.26658) genetic groups (*K*). We reconstructed a neighbor-joining maximum-likelihood tree with SNPhylo (Lee et al., 2014), based on genetic distances between all the individuals with 100 bootstraps to assess the reliability of the tree topology. We used plink (Purcell et al., 2007) to perform a principal component analysis (PCA) using 10 principal components.

### Coalescent simulations

We used coalescent simulations to test two different possible scenarios of divergence between the genetic groups observed in our ADMIXTURE and SNPhylo results. In the first scenario, *argyrosperma* descends from the *sororia* populations from Jalisco and the *sororia* populations from Southern Mexico coalesced with the *sororia* populations from Jalisco. In the second scenario, *argyrosperma* descends from the *sororia* populations from Southern Mexico and the *sororia* populations from Jalisco coalesced with the *sororia* populations from Southern Mexico. In addition, we also tested for each scenario of domestication whether divergence occurred in the presence of continuous gene flow, gene flow after secondary contact, or in the absence of gene flow.

We used Fastsimcoal 2 (Excoffier and Foll, 2011) to determine the parameters that maximize the composite likelihood of each model given the unfolded multidimensional SFS. The unfolded multidimensional SFS was obtained with DADI (Gutenkunst et al., 2009), using 17 *sororia* individuals of Jalisco, 27 *sororia* individuals of Southern Mexico, 109 *argyrosperma* individuals and 5 *C. moschata* individuals as an outgroup to unfold the SFS. We ran 100,000 simulations with 20 replicates for each model (two divergence scenarios and three gene flow scenarios) using the following settings: a parameter estimation by Maximum Likelihood with a stopping criterion of 0.001 difference between runs, a minimum SFS count of 1, a maximum of 40 loops to estimate the SFS parameters and a maximum of 200,000 simulations to estimate the SFS parameters. We also selected log-uniform priors for parameter estimations, setting times of divergence between 1000 and 200,000 generations (domestication times are expected to fall within this interval, given that the presumed most ancient evidence of human presence in America is 33,000 years old (Ardelean et al., 2020)), effective population sizes (*N*_e_) between 100 and 60,000 individuals and migration rates (m) between 0.0001 and 0.5. The 20 replicates of each model converged to similar likelihoods, indicating that the simulations performed well. After corroborating that all replicates converged to similar likelihoods, we combined all replicates and retained all outputs that were above the 95% of the likelihood distribution. We found that the Jalisco model of divergence with secondary contact had the lowest Akaike Information Criterion values for all the tested models.

### Tests to detect selective sweeps and introgression

We used BayeScEnv (de Villemereuil and Gaggiotti, 2015) to detect putative regions under selection that were differentiated between *sororia* and *argyrosperma*. For the “environmental” values used by BayeScEnv, we assigned each population as either wild (0) or domesticated (1). We ran two independent MCMC analyses with 20 initial pilot runs with a length of 10,000 generations and a main run with an initial burn-in of 100,000 generations and a subsequent sampling step for 100,000 generations sampling every 20 generations. We confirmed the convergence between both chains using the Gelman and Rubin statistic (Gelman and Rubin, 1992). The SNPs with *q-values* < 0.05 were regarded as candidate loci under selection.

The Mahalanobis distances implemented in PCAdapt (Luu et al., 2017) were used to detect candidate SNPs after controlling for the first two principal components in our dataset, which correspond to the subspecies and geographical differentiation observed among the populations (see Fig. S2). We performed Bonferroni corrections to adjust the *p-values* obtained from PCAdapt and the SNPs with Bonferroni corrected *p-values* < 0.05 were regarded as candidate loci under selection.

We performed an ABBA-BABA test using Dsuit (Malinsky et al., 2020) against the 11,498,421 whole-genome variants to evaluate signals of introgression or incomplete lineage sorting between *argyrosperma*, *sororia* and *C. moschata*, while using *C. okeechobeensis* subsp. *martinezii* as an outgroup. We calculated the *D*-statistic throughout the entire genome within 500 SNP windows with a step size of 250 SNPs. We also used the five tGBS data of *C. moschata*, as well as the whole-genome sequences of *C. moschata* and *C. okeechobeensis* subsp. *martinezii,* to define the ancestral state of each candidate locus and determine the direction of the selective signals.

We used snpEff (Cingolani et al., 2012) to associate the candidate loci found in both tests with the genome annotation of *argyrosperma* (Barrera-Redondo et al., 2019). The genes that could be unambiguously assigned to a candidate SNP (the SNP resided within the gene structure, including introns, exons and UTRs) were screened for a Gene Ontology enrichment analysis using topGO and the *weight01* algorithm (Alexa et al., 2006). We determined the significantly enriched biological functions by performing a Fisher’s exact test (*p*-value < 0.05).

### Accession numbers

The chromosome-level genome assemblies of *C. argyrosperma* subsp. *argyrosperma* and *C. argyrosperma* subsp. *sororia* are available at the Cucurbit Genomics Database (Zheng et al., 2019) and in the NCBI RefSeq database (accessions XXXXXXXXX). The raw sequencing reads of the *C. argyrosperma* subsp. *sororia, C. moschata* and *C. okeechobeensis* subsp. *martinezii* genomes are available in the NCBI Sequence Read Archive (accessions SRPXXXXXX- SRPXXXXXX). The raw sequencing reads of each individual sequenced by tGBS are available in the NCBI Sequence Read Archive (accessions SRPXXXXXX- SRPXXXXXX; table S3). The 13k SNP dataset is available in Dataset S1 and the 11,498,421 variant dataset is available in Dataset S2.

## Author Contributions

L.E.E., M.I.T., G.S-V. and J.B-R. designed the research; G.S-V. and G.C-M. collected the biological material; G.S-V extracted DNA for the tGBS data; G.C-M. and G.S-V. coordinated the sequencing procedures of the tGBS data; J.B-R. and X.A-D. extracted DNA for reference genome and coordinated its sequencing procedures; E.A-P. coordinated the administrative and lab work; J-B-R., J.A-L. and Y.T.G-G. performed the bioinformatic analyses; G.S-V., J.B-R., J.A-L., M.I.T., and L.E.E. analyzed the results; J.B-R., G.S-V. and L.E.E. drafted the manuscript; R.L-S. and L.E.E. obtained the funding. All authors revised the final version of the manuscript.

## Acknowledgments

We acknowledge the Doctorado en Ciencias Biomédicas for the support provided during the development of this project. We acknowledge the Doctorado en Ciencias Biológicas for the support provided during the development of this project. We thank Jodi Lynn Humann for her technical support while using the GenSAS web server. Special thanks to Laura Espinosa-Asuar and Silvia Barrientos for their technical support. We thank Rodrigo García Herrera, head of the Scientific Computing Department at LANCIS, Instituto de Ecología UNAM, for running the HTC infrastructure we used for the genome assembly and chromosome anchoring.

