## Supplementary information for "A wild *Cucurbita* genome reveals the role of structural variants and introgression in domestication"

#### This file includes:

**Supplementary Tables**
S1: Assembly metrics of *C. argyrosperma* subsp. *sororia* genome.
S2: Putative rearrangements between *argyrosperma* and *sororia.*
S3: Information of 192 individuals sequenced with tGBS.
S4: Genetic diversity of *C. argyrosperma* subspecies.
S5: Genetic diversity of each *C. argyrosperma* population.
S6: Introgression analysis using the ABBA-BABA test.
S7: Candidate genes under selection.
S8: GO enrichment of candidate genes.
S9: Structural variants with selective signals.

**Supplementary Figures**
S1: Gene synteny dot plots between *Cucurbita* genomes.
S2: Principal Component Analyses.
S3: LD decay in *C. argyrosperma*.
S4: ABBA-BABA test using sliding windows.

#### Supplementary Tables

**Table S1.** Assembly metrics of the genome of *Cucurbita argyrosperma* subsp. *sororia*.

| | |
|---|---|
| Assembly size | 253,587,946 bp |
| No. of contigs | 828 |
| Longest contig | 4,976,248 bp |
| N50 | 1,323,288 bp |
| L50 | 58 contigs |
| No. of contigs > 1 kbp | 828 (100.0%) |
| No. of contigs > 10 kbp | 810 (97.8%) |
| No. of contigs > 100 kbp | 292 (35.3%) |
| CG content | 36.56% |
| Illumina read coverage | 213x |
| PacBio read coverage | 75.4x |
| No. of genes | 31,452 |
| Average gene size | 3,269 bp |
| Complete BUSCOs | 92.80% |
| Fragmented BUSCOs | 1.20% |
| Missing BUSCOs | 6.00% |

**Table S2.** Putative rearrangements found between the genome assemblies of *C. argyrosperma* subsp. *sororia* (wild genome) and *C. argyrosperma* subsp. *argyrosperma* (domesticated genome) after aligning both genomes. Copy-number variants are considered gains or losses with respect to the domesticated genome.

| **Structural variants** | **Number of variants** | **Cumulative size of variants (bp)** | **Genes within variants** | **Enriched GO biological functions (*p*-value)** |
|---|---|---|---|---|
| Copy-gain variants | 878 | 4479959 | 152 | Regulation of transcription by RNA polymerase II (0.00017)<br>Pseudouridine synthesis (0.00027)<br>Cysteine biosynthetic process from serine (0.00142)<br>Protein transport by the Tat complex (0.00953)<br>Asparaginyl-tRNA aminoacylation (0.01898)<br>tRNA methylation (0.01898)<br>Lipid glycosylation (0.02833)<br>Nitrogen compound metabolic process (0.04653) |
| Copy-loss variants | 3381 | 15484147 | 610 | Transmembrane transport (6.3e-10)<br>Microtubule-based process (0.0015)<br>Meiotic chromosome segregation (0.0260)<br>Cell wall modification (0.0334)<br>7-methylguanosine mRNA capping (0.0388)<br>Protein lipoylation (0.0388)<br>mRNA methylation (0.0388) |
| Translocations | 1147 | 19604703 | 1297 | Protein dephosphorylation (0.002)<br>Intracellular sterol transport (0.002)<br>Molybdopterin cofactor biosynthetic process (0.002)<br>Golgi vesicle transport (0.014)<br>Protein peptidyl-prolyl isomerization (0.017)<br>Phenylalanyl-tRNA aminoacylation (0.027)<br>Protein-containing complex assembly (0.045) |
| Inversions | 46 | 1299618 | 46 | Glycolytic process (0.00014)<br>Isoleucine biosynthetic process (0.00624) |
| Unaligned regions in domesticated genome | 2479 | 18979604 | 149 | Cytoskeleton organization (0.0045)<br>NADP biosynthetic process (0.0182)<br>Tryptophan biosynthetic process (0.0212) |

| | | | | |
|---|---|---|---|---|
| | | | | Negative regulation of translation (0.0272)<br>Protein ubiquitination (0.0275)<br>Lysine biosynthetic process via diaminopimelate (0.0302)<br>Inositol phosphate dephosphorylation (0.0391)<br>Translational termination (0.0450) |
| Unaligned regions in wild genome | 3846 | 28437839 | 637 | Proteolysis (0.0005)<br>Tryptophan biosynthetic process (0.0012)<br>DNA replication initiation (0.0050)<br>Phosphorelay signal transduction system (0.0123)<br>ATP synthesis coupled proton transport (0.0130)<br>Negative regulation of DNA helicase activity (0.0134)<br>Glucosylceramide catabolic process (0.0266)<br>Sucrose biosynthetic process (0.0266)<br>Cytoskeleton organization (0.0307)<br>L-phenylalanine biosynthetic process (0.0460)<br>Cytochrome complex assembly (0.0460) |

**Table S3.** Information of 192 individuals sequenced using tGBS libraries, including population name, geographical coordinates and SRA accession. (within population names: W = wild, D = domesticated)

| Individual ID | Population name | Population number | Taxon | Latitude | Longitude | SRA accession |
|---|---|---|---|---|---|---|
| M_SON1 | Alamos, Sonora (Outgroup) | 0 | *Cucurbita moschata* (Outgroup) | 27.02694 | -108.93659 | XXXXXX |
| M_SON2 | Alamos, Sonora (Outgroup) | 0 | *Cucurbita moschata* (Outgroup) | 27.02694 | -108.93659 | XXXXXX |
| M_SON3 | Alamos, Sonora (Outgroup) | 0 | *Cucurbita moschata* (Outgroup) | 27.02694 | -108.93659 | XXXXXX |
| M_SON4 | Alamos, Sonora (Outgroup) | 0 | *Cucurbita moschata* (Outgroup) | 27.02694 | -108.93659 | XXXXXX |
| M_SON5 | Alamos, Sonora (Outgroup) | 0 | *Cucurbita moschata* (Outgroup) | 27.02694 | -108.93659 | XXXXXX |
| M_SON6 | Alamos, Sonora (Outgroup) | 0 | *Cucurbita moschata* (Outgroup) | 27.02694 | -108.93659 | XXXXXX |
| S_CHIS1 | Jiquipilas, Chiapas (W) | 1 | *Cucurbita argyrosperma* subsp. *sororia* | 16.597486 | -93.626878 | XXXXXX |
| S_CHIS2 | Jiquipilas, Chiapas (W) | 1 | *Cucurbita argyrosperma* subsp. *sororia* | 16.597486 | -93.626878 | XXXXXX |
| S_CHIS3 | Jiquipilas, Chiapas (W) | 1 | *Cucurbita argyrosperma* subsp. *sororia* | 16.597486 | -93.626878 | XXXXXX |
| S_CHIS4 | Jiquipilas, Chiapas (W) | 1 | *Cucurbita argyrosperma* subsp. *sororia* | 16.597486 | -93.626878 | XXXXXX |
| S_CHIS5 | Jiquipilas, Chiapas (W) | 1 | *Cucurbita argyrosperma* subsp. *sororia* | 16.597486 | -93.626878 | XXXXXX |
| S_CHIS6 | Jiquipilas, Chiapas (W) | 1 | *Cucurbita argyrosperma* subsp. *sororia* | 16.597486 | -93.626878 | XXXXXX |
| S_CHIS7 | Jiquipilas, Chiapas (W) | 1 | *Cucurbita argyrosperma* subsp. *sororia* | 16.597486 | -93.626878 | XXXXXX |
| S_CHIS8 | Jiquipilas, Chiapas (W) | 1 | *Cucurbita argyrosperma* subsp. *sororia* | 16.597486 | -93.626878 | XXXXXX |
| S_GRO1 | Ometepec, Guerrero (W) | 2 | *Cucurbita argyrosperma* subsp. *sororia* | 16.688139 | -98.406319 | XXXXXX |
| S_GRO2 | Ometepec, Guerrero (W) | 2 | *Cucurbita argyrosperma* subsp. *sororia* | 16.688139 | -98.406319 | XXXXXX |
| S_GRO3 | Ometepec, Guerrero (W) | 2 | *Cucurbita argyrosperma* subsp. *sororia* | 16.688139 | -98.406319 | XXXXXX |
| S_GRO4 | Ometepec, Guerrero (W) | 2 | *Cucurbita argyrosperma* subsp. *sororia* | 16.688139 | -98.406319 | XXXXXX |
| S_GRO5 | Ometepec, Guerrero (W) | 2 | *Cucurbita argyrosperma* subsp. *sororia* | 16.688139 | -98.406319 | XXXXXX |

| | | | | | | |
|---|---|---|---|---|---|---|
| S_OAX1 | Puerto Escondido, Oaxaca (W) | 3 | *Cucurbita argyrosperma* subsp. *sororia* | 15.918225 | -97.076308 | XXXXXX |
| S_OAX2 | Puerto Escondido, Oaxaca (W) | 3 | *Cucurbita argyrosperma* subsp. *sororia* | 15.918225 | -97.076308 | XXXXXX |
| S_OAX3 | Puerto Escondido, Oaxaca (W) | 3 | *Cucurbita argyrosperma* subsp. *sororia* | 15.918225 | -97.076308 | XXXXXX |
| S_OAX4 | Puerto Escondido, Oaxaca (W) | 3 | *Cucurbita argyrosperma* subsp. *sororia* | 15.918225 | -97.076308 | XXXXXX |
| S_OAX5 | Puerto Escondido, Oaxaca (W) | 3 | *Cucurbita argyrosperma* subsp. *sororia* | 15.918225 | -97.076308 | XXXXXX |
| S_OAX6 | Puerto Escondido, Oaxaca (W) | 3 | *Cucurbita argyrosperma* subsp. *sororia* | 15.918225 | -97.076308 | XXXXXX |
| S_OAX7 | Puerto Escondido, Oaxaca (W) | 3 | *Cucurbita argyrosperma* subsp. *sororia* | 15.918225 | -97.076308 | XXXXXX |
| S_OAX8 | Puerto Escondido, Oaxaca (W) | 3 | *Cucurbita argyrosperma* subsp. *sororia* | 15.918225 | -97.076308 | XXXXXX |
| S_OAX9 | Puerto Escondido, Oaxaca (W) | 3 | *Cucurbita argyrosperma* subsp. *sororia* | 15.918225 | -97.076308 | XXXXXX |
| S_OAX10 | Puerto Escondido, Oaxaca (W) | 3 | *Cucurbita argyrosperma* subsp. *sororia* | 15.918225 | -97.076308 | XXXXXX |
| S_OAX11 | Puerto Escondido, Oaxaca (W) | 3 | *Cucurbita argyrosperma* subsp. *sororia* | 15.9528333 | -97.0772778 | XXXXXX |

| | | | | | | |
|---|---|---|---|---|---|---|
| S_OAX12 | Puerto Escondido, Oaxaca (W) | 3 | *Cucurbita argyrosperma* subsp. *sororia* | 15.9528333 | -97.0772778 | XXXXXX |
| S_OAX13 | Puerto Escondido, Oaxaca (W) | 3 | *Cucurbita argyrosperma* subsp. *sororia* | 15.9528333 | -97.0772778 | XXXXXX |
| S_OAX14 | Puerto Escondido, Oaxaca (W) | 3 | *Cucurbita argyrosperma* subsp. *sororia* | 15.9528333 | -97.0772778 | XXXXXX |
| S_JAL1 | Jalisco (W) | 4 | *Cucurbita argyrosperma* subsp. *sororia* | 19.682 | -104.333278 | XXXXXX |
| S_JAL2 | Jalisco (W) | 4 | *Cucurbita argyrosperma* subsp. *sororia* | 19.682 | -104.333278 | XXXXXX |
| S_JAL3 | Jalisco (W) | 4 | *Cucurbita argyrosperma* subsp. *sororia* | 19.7019583 | -104.204314 | XXXXXX |
| S_JAL4 | Jalisco (W) | 4 | *Cucurbita argyrosperma* subsp. *sororia* | 19.9005833 | -104.160222 | XXXXXX |
| S_JAL5 | Jalisco (W) | 4 | *Cucurbita argyrosperma* subsp. *sororia* | 19.9005833 | -104.160222 | XXXXXX |
| S_JAL6 | Jalisco (W) | 4 | *Cucurbita argyrosperma* subsp. *sororia* | 19.9005833 | -104.160222 | XXXXXX |
| S_JAL7 | Jalisco (W) | 4 | *Cucurbita argyrosperma* subsp. *sororia* | 19.9005833 | -104.160222 | XXXXXX |
| S_JAL8 | Jalisco (W) | 4 | *Cucurbita argyrosperma* subsp. *sororia* | 19.6997222 | -104.203056 | XXXXXX |
| S_JAL9 | Jalisco (W) | 4 | *Cucurbita argyrosperma* subsp. *sororia* | 19.6997222 | -104.203056 | XXXXXX |
| S_JAL10 | Jalisco (W) | 4 | *Cucurbita argyrosperma* subsp. *sororia* | 19.6997222 | -104.203056 | XXXXXX |
| S_JAL11 | Jalisco (W) | 4 | *Cucurbita argyrosperma* subsp. *sororia* | 19.8753889 | -104.072333 | XXXXXX |
| S_JAL12 | Jalisco (W) | 4 | *Cucurbita argyrosperma* subsp. *sororia* | 19.9123056 | -104.116833 | XXXXXX |
| S_JAL13 | Jalisco (W) | 4 | *Cucurbita argyrosperma* subsp. *sororia* | 19.9123056 | -104.116833 | XXXXXX |
| S_JAL14 | Jalisco (W) | 4 | *Cucurbita argyrosperma* subsp. *sororia* | 19.9123056 | -104.116833 | XXXXXX |
| S_JAL15 | Jalisco (W) | 4 | *Cucurbita argyrosperma* subsp. *sororia* | 19.9123056 | -104.116833 | XXXXXX |
| S_JAL16 | Jalisco (W) | 4 | *Cucurbita argyrosperma* subsp. *sororia* | 19.9600833 | -104.037472 | XXXXXX |
| S_JAL17 | Jalisco (W) | 4 | *Cucurbita argyrosperma* subsp. *sororia* | 19.9574722 | -103.988389 | XXXXXX |

| | | | | | | |
|---|---|---|---|---|---|---|
| S_JAL18 | Jalisco (W) | 4 | *Cucurbita argyrosperma* subsp. *sororia* | 19.8753889 | -104.072333 | XXXXXX |
| S_JAL19 | Jalisco (W) | 4 | *Cucurbita argyrosperma* subsp. *sororia* | 19.8356111 | -104.081417 | XXXXXX |
| S_JAL20 | Jalisco (W) | 4 | *Cucurbita argyrosperma* subsp. *sororia* | 19.8356111 | -104.081417 | XXXXXX |
| S_JAL21 | Jalisco (W) | 4 | *Cucurbita argyrosperma* subsp. *sororia* | 19.6909722 | -104.360833 | XXXXXX |
| S_JAL22 | Jalisco (W) | 4 | *Cucurbita argyrosperma* subsp. *sororia* | 19.6909722 | -104.360833 | XXXXXX |
| S_JAL23 | Jalisco (W) | 4 | *Cucurbita argyrosperma* subsp. *sororia* | 19.6909722 | -104.360833 | XXXXXX |
| A_TLAP1 | Tlapehuala, Guerrero (D) | 5 | *Cucurbita argyrosperma* subsp. *argyrosperma* | 18.2416667 | -100.534722 | XXXXXX |
| A_TLAP2 | Tlapehuala, Guerrero (D) | 5 | *Cucurbita argyrosperma* subsp. *argyrosperma* | 18.2416667 | -100.534722 | XXXXXX |
| A_TLAP3 | Tlapehuala, Guerrero (D) | 5 | *Cucurbita argyrosperma* subsp. *argyrosperma* | 18.2416667 | -100.534722 | XXXXXX |
| A_TLAP4 | Tlapehuala, Guerrero (D) | 5 | *Cucurbita argyrosperma* subsp. *argyrosperma* | 18.2416667 | -100.534722 | XXXXXX |
| A_TLAP5 | Tlapehuala, Guerrero (D) | 5 | *Cucurbita argyrosperma* subsp. *argyrosperma* | 18.2416667 | -100.534722 | XXXXXX |
| A_TLAP6 | Tlapehuala, Guerrero (D) | 5 | *Cucurbita argyrosperma* subsp. *argyrosperma* | 18.2416667 | -100.534722 | XXXXXX |
| A_TLAP7 | Tlapehuala, Guerrero (D) | 5 | *Cucurbita argyrosperma* subsp. *argyrosperma* | 18.2416667 | -100.534722 | XXXXXX |
| A_TLAP8 | Tlapehuala, Guerrero (D) | 5 | *Cucurbita argyrosperma* subsp. *argyrosperma* | 18.2416667 | -100.534722 | XXXXXX |
| A_TLAP9 | Tlapehuala, Guerrero (D) | 5 | *Cucurbita argyrosperma* subsp. *argyrosperma* | 18.2416667 | -100.534722 | XXXXXX |
| A_TLAP10 | Tlapehuala, Guerrero (D) | 5 | *Cucurbita argyrosperma* subsp. *argyrosperma* | 18.2416667 | -100.534722 | XXXXXX |
| A_JAL1 | Jalisco (D) | 6 | *Cucurbita argyrosperma* subsp. *argyrosperma* | 19.7019583 | -104.204314 | XXXXXX |
| A_JAL2 | Jalisco (D) | 6 | *Cucurbita argyrosperma* subsp. *argyrosperma* | 19.7019583 | -104.204314 | XXXXXX |
| A_JAL3 | Jalisco (D) | 6 | *Cucurbita argyrosperma* subsp. *argyrosperma* | 19.6997222 | -104.203056 | XXXXXX |
| A_JAL4 | Jalisco (D) | 6 | *Cucurbita argyrosperma* subsp. *argyrosperma* | 19.6997222 | -104.203056 | XXXXXX |
| A_JAL5 | Jalisco (D) | 6 | *Cucurbita argyrosperma* subsp. *argyrosperma* | 19.8753889 | -104.072333 | XXXXXX |
| A_JAL6 | Jalisco (D) | 6 | *Cucurbita argyrosperma* subsp. *argyrosperma* | 19.8753889 | -104.072333 | XXXXXX |
| A_JAL7 | Jalisco (D) | 6 | *Cucurbita argyrosperma* subsp. *argyrosperma* | 19.9600833 | -104.037472 | XXXXXX |

| | | | | | | |
|---|---|---|---|---|---|---|
| A_JAL8 | Jalisco (D) | 6 | *Cucurbita argyrosperma* subsp. *argyrosperma* | 19.8714722 | -104.217333 | XXXXXX |
| A_JAL9 | Jalisco (D) | 6 | *Cucurbita argyrosperma* subsp. *argyrosperma* | 19.8714722 | -104.217333 | XXXXXX |
| A_JAL10 | Jalisco (D) | 6 | *Cucurbita argyrosperma* subsp. *argyrosperma* | 19.9574722 | -103.988389 | XXXXXX |
| A_JAL11 | Jalisco (D) | 6 | *Cucurbita argyrosperma* subsp. *argyrosperma* | 19.8356111 | -104.081417 | XXXXXX |
| A_JAL12 | Jalisco (D) | 6 | *Cucurbita argyrosperma* subsp. *argyrosperma* | 19.8356111 | -104.081417 | XXXXXX |
| A_JAL13 | Jalisco (D) | 6 | *Cucurbita argyrosperma* subsp. *argyrosperma* | 19.8356111 | -104.081417 | XXXXXX |
| A_JAL14 | Jalisco (D) | 6 | *Cucurbita argyrosperma* subsp. *argyrosperma* | 19.8356111 | -104.081417 | XXXXXX |
| A_JAL15 | Jalisco (D) | 6 | *Cucurbita argyrosperma* subsp. *argyrosperma* | 19.8356111 | -104.081417 | XXXXXX |
| A_JAL16 | Jalisco (D) | 6 | *Cucurbita argyrosperma* subsp. *argyrosperma* | 19.8356111 | -104.081417 | XXXXXX |
| A_JAL17 | Jalisco (D) | 6 | *Cucurbita argyrosperma* subsp. *argyrosperma* | 19.8356111 | -104.081417 | XXXXXX |
| A_BAD1 | Badiraguato, Sinaloa (D) | 7 | *Cucurbita argyrosperma* subsp. *argyrosperma* | 25.359173 | -107.558408 | XXXXXX |
| A_MTP1 | Matlalapa, Guerrero (D) | 8 | *Cucurbita argyrosperma* subsp. *argyrosperma* | 17.5971639 | -99.4579917 | XXXXXX |
| A_MTP2 | Matlalapa, Guerrero (D) | 8 | *Cucurbita argyrosperma* subsp. *argyrosperma* | 17.5971639 | -99.4579917 | XXXXXX |
| A_MTP3 | Matlalapa, Guerrero (D) | 8 | *Cucurbita argyrosperma* subsp. *argyrosperma* | 17.5971639 | -99.4579917 | XXXXXX |
| A_MTP4 | Matlalapa, Guerrero (D) | 8 | *Cucurbita argyrosperma* subsp. *argyrosperma* | 17.5971639 | -99.4579917 | XXXXXX |
| A_MTP5 | Matlalapa, Guerrero (D) | 8 | *Cucurbita argyrosperma* subsp. *argyrosperma* | 17.5971639 | -99.4579917 | XXXXXX |
| A_MTP6 | Matlalapa, Guerrero (D) | 8 | *Cucurbita argyrosperma* subsp. *argyrosperma* | 17.5971639 | -99.4579917 | XXXXXX |
| A_MTP7 | Matlalapa, Guerrero (D) | 8 | *Cucurbita argyrosperma* subsp. *argyrosperma* | 17.5971639 | -99.4579917 | XXXXXX |
| A_SAH1 | Sahuayo, Michoacán (D) | 9 | *Cucurbita argyrosperma* subsp. *argyrosperma* | 20.0587194 | -102.716233 | XXXXXX |
| A_SAH2 | Sahuayo, Michoacán (D) | 9 | *Cucurbita argyrosperma* subsp. *argyrosperma* | 20.0587194 | -102.716233 | XXXXXX |
| A_SAH3 | Sahuayo, Michoacán (D) | 9 | *Cucurbita argyrosperma* subsp. *argyrosperma* | 20.0587194 | -102.716233 | XXXXXX |
| A_SAH4 | Sahuayo, Michoacán (D) | 9 | *Cucurbita argyrosperma* subsp. *argyrosperma* | 20.0587194 | -102.716233 | XXXXXX |
| A_SAH5 | Sahuayo, Michoacán (D) | 9 | *Cucurbita argyrosperma* subsp. *argyrosperma* | 20.0587194 | -102.716233 | XXXXXX |

| | | | | | | |
|---|---|---|---|---|---|---|
| A_SAH6 | Sahuayo, Michoacán (D) | 9 | *Cucurbita argyrosperma* subsp. *argyrosperma* | 20.0587194 | -102.716233 | XXXXXX |
| A_SAH7 | Sahuayo, Michoacán (D) | 9 | *Cucurbita argyrosperma* subsp. *argyrosperma* | 20.0587194 | -102.716233 | XXXXXX |
| A_SAH8 | Sahuayo, Michoacán (D) | 9 | *Cucurbita argyrosperma* subsp. *argyrosperma* | 20.0587194 | -102.716233 | XXXXXX |
| A_SAH9 | Sahuayo, Michoacán (D) | 9 | *Cucurbita argyrosperma* subsp. *argyrosperma* | 20.0587194 | -102.716233 | XXXXXX |
| A_SAH10 | Sahuayo, Michoacán (D) | 9 | *Cucurbita argyrosperma* subsp. *argyrosperma* | 20.0587194 | -102.716233 | XXXXXX |
| A_SAL1 | Salamanca, Guanajuato (D) | 10 | *Cucurbita argyrosperma* subsp. *argyrosperma* | 20.5205778 | -101.190992 | XXXXXX |
| A_SJI1 | San José Iturbide, Guanajuato (D) | 10 | *Cucurbita argyrosperma* subsp. *argyrosperma* | 20.9988889 | -100.385 | XXXXXX |
| A_NAY1 | Tepic, Nayarit (D) | 11 | *Cucurbita argyrosperma* subsp. *argyrosperma* | 21.519956 | -104.893423 | XXXXXX |
| A_NAY2 | Tepic, Nayarit (D) | 11 | *Cucurbita argyrosperma* subsp. *argyrosperma* | 21.519956 | -104.893423 | XXXXXX |
| A_NAY3 | Tepic, Nayarit (D) | 11 | *Cucurbita argyrosperma* subsp. *argyrosperma* | 21.519956 | -104.893423 | XXXXXX |
| A_NAY4 | Tepic, Nayarit (D) | 11 | *Cucurbita argyrosperma* subsp. *argyrosperma* | 21.519956 | -104.893423 | XXXXXX |
| F_PLAT1 | El Platanar, Sinaloa (feral) | 12 | feral individual | 24.0303667 | -106.432561 | XXXXXX |
| F_PLAT2 | El Platanar, Sinaloa (feral) | 12 | feral individual | 24.0303667 | -106.432561 | XXXXXX |
| F_PLAT3 | El Platanar, Sinaloa (feral) | 12 | feral individual | 24.0303667 | -106.432561 | XXXXXX |
| F_PLAT4 | El Platanar, Sinaloa (feral) | 12 | feral individual | 24.0303667 | -106.432561 | XXXXXX |
| F_PLAT5 | El Platanar, Sinaloa (feral) | 12 | feral individual | 24.0303667 | -106.432561 | XXXXXX |
| F_PLAT6 | El Platanar, Sinaloa (feral) | 12 | feral individual | 24.0303667 | -106.432561 | XXXXXX |
| F_PLAT7 | El Platanar, Sinaloa (feral) | 12 | feral individual | 24.0303667 | -106.432561 | XXXXXX |
| F_CUL1 | Culiacán, Sinaloa (feral) | 13 | feral individual | 24.817335 | -107.416667 | XXXXXX |
| F_CUL2 | Culiacán, Sinaloa (feral) | 13 | feral individual | 24.817335 | -107.416667 | XXXXXX |
| F_CUL3 | Culiacán, Sinaloa (feral) | 13 | feral individual | 24.817335 | -107.416667 | XXXXXX |
| F_CUL4 | Culiacán, Sinaloa (feral) | 13 | feral individual | 24.817335 | -107.416667 | XXXXXX |

| | | | | | | |
|---|---|---|---|---|---|---|
| F_CUL5 | Culiacán, Sinaloa (feral) | 13 | feral individual | 24.817335 | -107.416667 | XXXXXX |
| A_CHOI1 | Choix, Sinaloa (D) | 14 | *Cucurbita argyrosperma* subsp. *argyrosperma* | 26.5967833 | -108.335581 | XXXXXX |
| A_CHOI2 | Choix, Sinaloa (D) | 14 | *Cucurbita argyrosperma* subsp. *argyrosperma* | 26.5967833 | -108.335581 | XXXXXX |
| A_CHOI3 | Choix, Sinaloa (D) | 14 | *Cucurbita argyrosperma* subsp. *argyrosperma* | 26.5967833 | -108.335581 | XXXXXX |
| A_CHOI4 | Choix, Sinaloa (D) | 14 | *Cucurbita argyrosperma* subsp. *argyrosperma* | 26.5967833 | -108.335581 | XXXXXX |
| A_CHOI5 | Choix, Sinaloa (D) | 14 | *Cucurbita argyrosperma* subsp. *argyrosperma* | 26.5967833 | -108.335581 | XXXXXX |
| A_YEC1 | Yecora, Sonora (D) | 15 | *Cucurbita argyrosperma* subsp. *argyrosperma* | 28.3720417 | -108.926986 | XXXXXX |
| A_YEC2 | Yecora, Sonora (D) | 15 | *Cucurbita argyrosperma* subsp. *argyrosperma* | 28.3720417 | -108.926986 | XXXXXX |
| A_YEC3 | Yecora, Sonora (D) | 15 | *Cucurbita argyrosperma* subsp. *argyrosperma* | 28.3720417 | -108.926986 | XXXXXX |
| A_YEC4 | Yecora, Sonora (D) | 15 | *Cucurbita argyrosperma* subsp. *argyrosperma* | 28.3720417 | -108.926986 | XXXXXX |
| A_YEC5 | Yecora, Sonora (D) | 15 | *Cucurbita argyrosperma* subsp. *argyrosperma* | 28.3720417 | -108.926986 | XXXXXX |
| A_YEC6 | Yecora, Sonora (D) | 15 | *Cucurbita argyrosperma* subsp. *argyrosperma* | 28.3720417 | -108.926986 | XXXXXX |
| F_ONAV1 | Onavas, Sonora (feral) | 16 | feral individual | 28.533333 | -109.583333 | XXXXXX |
| F_ONAV2 | Onavas, Sonora (feral) | 16 | feral individual | 28.533333 | -109.583333 | XXXXXX |
| F_ONAV3 | Onavas, Sonora (feral) | 16 | feral individual | 28.533333 | -109.583333 | XXXXXX |
| F_ONAV4 | Onavas, Sonora (feral) | 16 | feral individual | 28.533333 | -109.583333 | XXXXXX |
| F_ONAV5 | Onavas, Sonora (feral) | 16 | feral individual | 28.533333 | -109.583333 | XXXXXX |
| F_ONAV6 | Onavas, Sonora (feral) | 16 | feral individual | 28.533333 | -109.583333 | XXXXXX |
| F_ONAV7 | Onavas, Sonora (feral) | 16 | feral individual | 28.533333 | -109.583333 | XXXXXX |
| A_DGO1 | Durango, Durango (D) | 17 | *Cucurbita argyrosperma* subsp. *argyrosperma* | 24.066667 | -104.583333 | XXXXXX |
| A_TEH1 | Tehuantepec, Oaxaca (D) | 18 | *Cucurbita argyrosperma* subsp. *argyrosperma* | 16.3328306 | -95.2330361 | XXXXXX |
| A_TEH2 | Tehuantepec, Oaxaca (D) | 18 | *Cucurbita argyrosperma* subsp. *argyrosperma* | 16.3328306 | -95.2330361 | XXXXXX |
| A_TEH3 | Tehuantepec, Oaxaca (D) | 18 | *Cucurbita argyrosperma* subsp. *argyrosperma* | 16.3328306 | -95.2330361 | XXXXXX |

| | | | | | | |
|---|---|---|---|---|---|---|
| A_TEH4 | Tehuantepec, Oaxaca (D) | 18 | *Cucurbita argyrosperma* subsp. *argyrosperma* | 16.3328306 | -95.2330361 | XXXXXX |
| A_TEH5 | Tehuantepec, Oaxaca (D) | 18 | *Cucurbita argyrosperma* subsp. *argyrosperma* | 16.3328306 | -95.2330361 | XXXXXX |
| A_TEH6 | Tehuantepec, Oaxaca (D) | 18 | *Cucurbita argyrosperma* subsp. *argyrosperma* | 16.3328306 | -95.2330361 | XXXXXX |
| A_TEH7 | Tehuantepec, Oaxaca (D) | 18 | *Cucurbita argyrosperma* subsp. *argyrosperma* | 16.3328306 | -95.2330361 | XXXXXX |
| A_ONAV1 | Onavas, Sonora (D) | 19 | *Cucurbita argyrosperma* subsp. *argyrosperma* | 28.533333 | -109.583333 | XXXXXX |
| A_ONAV2 | Onavas, Sonora (D) | 19 | *Cucurbita argyrosperma* subsp. *argyrosperma* | 28.533333 | -109.583333 | XXXXXX |
| A_ONAV3 | Onavas, Sonora (D) | 19 | *Cucurbita argyrosperma* subsp. *argyrosperma* | 28.533333 | -109.583333 | XXXXXX |
| A_ONAV4 | Onavas, Sonora (D) | 19 | *Cucurbita argyrosperma* subsp. *argyrosperma* | 28.533333 | -109.583333 | XXXXXX |
| A_ONAV5 | Onavas, Sonora (D) | 19 | *Cucurbita argyrosperma* subsp. *argyrosperma* | 28.533333 | -109.583333 | XXXXXX |
| A_ONAV6 | Onavas, Sonora (D) | 19 | *Cucurbita argyrosperma* subsp. *argyrosperma* | 28.533333 | -109.583333 | XXXXXX |
| A_VER1 | Tihuatlán, Veracruz (D) | 20 | *Cucurbita argyrosperma* subsp. *argyrosperma* | 20.7200667 | -97.5395028 | XXXXXX |
| A_VER2 | Tihuatlán, Veracruz (D) | 20 | *Cucurbita argyrosperma* subsp. *argyrosperma* | 20.7200667 | -97.5395028 | XXXXXX |
| A_VER3 | Tihuatlán, Veracruz (D) | 20 | *Cucurbita argyrosperma* subsp. *argyrosperma* | 20.7200667 | -97.5395028 | XXXXXX |
| A_VER4 | Tihuatlán, Veracruz (D) | 20 | *Cucurbita argyrosperma* subsp. *argyrosperma* | 20.7200667 | -97.5395028 | XXXXXX |
| A_VER5 | Tihuatlán, Veracruz (D) | 20 | *Cucurbita argyrosperma* subsp. *argyrosperma* | 20.7200667 | -97.5395028 | XXXXXX |
| A_PAL1 | Palenque, Chiapas (D) | 21 | *Cucurbita argyrosperma* subsp. *argyrosperma* | 17.5128139 | -91.9877611 | XXXXXX |
| A_PAL2 | Palenque, Chiapas (D) | 21 | *Cucurbita argyrosperma* subsp. *argyrosperma* | 17.5128139 | -91.9877611 | XXXXXX |
| A_PAL3 | Palenque, Chiapas (D) | 21 | *Cucurbita argyrosperma* subsp. *argyrosperma* | 17.5128139 | -91.9877611 | XXXXXX |
| A_PAL4 | Palenque, Chiapas (D) | 21 | *Cucurbita argyrosperma* subsp. *argyrosperma* | 17.5128139 | -91.9877611 | XXXXXX |
| A_PAL5 | Palenque, Chiapas (D) | 21 | *Cucurbita argyrosperma* subsp. *argyrosperma* | 17.5128139 | -91.9877611 | XXXXXX |
| A_PAL6 | Palenque, Chiapas (D) | 21 | *Cucurbita argyrosperma* subsp. *argyrosperma* | 17.5128139 | -91.9877611 | XXXXXX |
| A_SLP1 | Tanquián, San Luis Potosí (D) | 22 | *Cucurbita argyrosperma* subsp. *argyrosperma* | 22.1154861 | -101.009331 | XXXXXX |

| | | | | | | |
|---|---|---|---|---|---|---|
| A_SLP2 | Tanquián, San Luis Potosí (D) | 22 | *Cucurbita argyrosperma* subsp. *argyrosperma* | 22.1154861 | -101.009331 | XXXXXX |
| A_SLP3 | Tanquián, San Luis Potosí (D) | 22 | *Cucurbita argyrosperma* subsp. *argyrosperma* | 22.1154861 | -101.009331 | XXXXXX |
| A_SLP4 | Tanquián, San Luis Potosí (D) | 22 | *Cucurbita argyrosperma* subsp. *argyrosperma* | 22.1154861 | -101.009331 | XXXXXX |
| A_SLP5 | Tanquián, San Luis Potosí (D) | 22 | *Cucurbita argyrosperma* subsp. *argyrosperma* | 22.1154861 | -101.009331 | XXXXXX |
| A_SLP6 | Tanquián, San Luis Potosí (D) | 22 | *Cucurbita argyrosperma* subsp. *argyrosperma* | 22.1154861 | -101.009331 | XXXXXX |
| A_CHAMP1 | Champotón, Campeche (D) | 23 | *Cucurbita argyrosperma* subsp. *argyrosperma* | 19.5011556 | -90.4613778 | XXXXXX |
| A_CHAMP2 | Champotón, Campeche (D) | 23 | *Cucurbita argyrosperma* subsp. *argyrosperma* | 19.5011556 | -90.4613778 | XXXXXX |
| A_CHAMP3 | Champotón, Campeche (D) | 23 | *Cucurbita argyrosperma* subsp. *argyrosperma* | 19.5011556 | -90.4613778 | XXXXXX |
| A_CHAMP4 | Champotón, Campeche (D) | 23 | *Cucurbita argyrosperma* subsp. *argyrosperma* | 19.5011556 | -90.4613778 | XXXXXX |
| A_CHAMP5 | Champotón, Campeche (D) | 23 | *Cucurbita argyrosperma* subsp. *argyrosperma* | 19.5011556 | -90.4613778 | XXXXXX |
| A_MIXT1 | Mixtepec, Oaxaca (D) | 24 | *Cucurbita argyrosperma* subsp. *argyrosperma* | 15.95875 | -97.0849167 | XXXXXX |
| A_MIXT2 | Mixtepec, Oaxaca (D) | 24 | *Cucurbita argyrosperma* subsp. *argyrosperma* | 15.95875 | -97.0849167 | XXXXXX |
| A_MIXT3 | Mixtepec, Oaxaca (D) | 24 | *Cucurbita argyrosperma* subsp. *argyrosperma* | 15.95875 | -97.0849167 | XXXXXX |
| A_MIXT4 | Mixtepec, Oaxaca (D) | 24 | *Cucurbita argyrosperma* subsp. *argyrosperma* | 15.95875 | -97.0849167 | XXXXXX |
| A_MIXT5 | Mixtepec, Oaxaca (D) | 24 | *Cucurbita argyrosperma* subsp. *argyrosperma* | 15.95875 | -97.0849167 | XXXXXX |
| A_MIXT6 | Mixtepec, Oaxaca (D) | 24 | *Cucurbita argyrosperma* subsp. *argyrosperma* | 15.95875 | -97.0849167 | XXXXXX |
| A_EK1 | Ek Balam, Yucatan (D) | 25 | *Cucurbita argyrosperma* subsp. *argyrosperma* | 20.9166667 | -87.9166667 | XXXXXX |
| A_EK2 | Ek Balam, Yucatan (D) | 25 | *Cucurbita argyrosperma* subsp. *argyrosperma* | 20.9166667 | -87.9166667 | XXXXXX |

| | | | | | | |
|---|---|---|---|---|---|---|
| A_EK3 | Ek Balam, Yucatan (D) | 25 | *Cucurbita argyrosperma* subsp. *argyrosperma* | 20.9166667 | -87.9166667 | XXXXXX |
| A_EK4 | Ek Balam, Yucatan (D) | 25 | *Cucurbita argyrosperma* subsp. *argyrosperma* | 20.9166667 | -87.9166667 | XXXXXX |
| A_EK5 | Ek Balam, Yucatan (D) | 25 | *Cucurbita argyrosperma* subsp. *argyrosperma* | 20.9166667 | -87.9166667 | XXXXXX |
| A_EK6 | Ek Balam, Yucatan (D) | 25 | *Cucurbita argyrosperma* subsp. *argyrosperma* | 20.9166667 | -87.9166667 | XXXXXX |
| A_CHAN1 | Chan Santa Cruz, Quintana Roo (D) | 26 | *Cucurbita argyrosperma* subsp. *argyrosperma* | 19.3670639 | -88.3328917 | XXXXXX |
| A_CHAN2 | Chan Santa Cruz, Quintana Roo (D) | 26 | *Cucurbita argyrosperma* subsp. *argyrosperma* | 19.3670639 | -88.3328917 | XXXXXX |
| A_CHAN3 | Chan Santa Cruz, Quintana Roo (D) | 26 | *Cucurbita argyrosperma* subsp. *argyrosperma* | 19.3670639 | -88.3328917 | XXXXXX |
| A_CHAN4 | Chan Santa Cruz, Quintana Roo (D) | 26 | *Cucurbita argyrosperma* subsp. *argyrosperma* | 19.3670639 | -88.3328917 | XXXXXX |
| A_CHAN5 | Chan Santa Cruz, Quintana Roo (D) | 26 | *Cucurbita argyrosperma* subsp. *argyrosperma* | 19.3670639 | -88.3328917 | XXXXXX |
| A_CHAN6 | Chan Santa Cruz, Quintana Roo (D) | 26 | *Cucurbita argyrosperma* subsp. *argyrosperma* | 19.3670639 | -88.3328917 | XXXXXX |
| A_CHAN7 | Chan Santa Cruz, Quintana Roo (D) | 26 | *Cucurbita argyrosperma* subsp. *argyrosperma* | 19.3670639 | -88.3328917 | XXXXXX |

**Table S4.** Average genetic diversity of the wild, domesticated and feral populations of *Cucurbita argyrosperma* using 2,861 unlinked SNPs predicted with the domesticated (D) reference genome, and 1,771 unlinked SNPs predicted with the wild (W) reference genome.

| **Taxon** | $N_{ind}$ | $N_{pop}$ | $H_O$ **(Var)** | | $H_E$ **(Var)** | | $\pi$ **(Var)** | | $F_{IS}$ **(Var)** | |
|---|---|---|---|---|---|---|---|---|---|---|
| | | | **D** | **W** | **D** | **W** | **D** | **W** | **D** | **W** |
| *Cucurbita argyrosperma* subsp. *sororia* | 44 | 4 | 0.098 (0.017) | 0.089 (0.016) | 0.096 (0.014) | 0.094 (0.015) | 0.098 (0.015) | 0.096 (0.015) | 0.011 (0.027) | 0.044 (0.037) |
| *Cucurbita argyrosperma* subsp. *argyrosperma* | 109 | 19 | 0.094 (0.012) | 0.095 (0.015) | 0.094 (0.010) | 0.097 (0.012) | 0.095 (0.010) | 0.098 (0.012) | 0.034 (0.030) | 0.067 (0.038) |
| feral populations | 14 | 3 | 0.102 (0.032) | 0.102 (0.035) | 0.088 (0.019) | 0.089 (0.022) | 0.094 (0.023) | 0.096 (0.026) | -0.015 (0.022) | -0.011 (0.026) |
| *Cucurbita moschata* (outgroup) | 5 | 1 | 0.077 (0.042) | 0.080 (0.047) | 0.058 (0.019) | 0.058 (0.021) | 0.068 (0.029) | 0.069 (0.032) | -0.017 (0.017) | -0.020 (0.019) |

($N_{ind}$ = number of individuals, $N_{pop}$ = number of populations, $H_O$ = observed heterozygosity, $H_E$ = expected heterozygosity, $\pi$ = nucleotide diversity, $F_{IS}$ = inbreeding coefficient, Var = variance)

**Table S5.** Genetic diversity of each wild, domesticated and feral population of *Cucurbita argyrosperma* using 2,861 unlinked SNPs (r2 < 0.25, MAF > 1%).

| Population name | Population number | *N* | $H_O$ (Var) | $H_E$ (Var) | $\pi$ (Var) | $F_{IS}$ (Var) |
|---|---|---|---|---|---|---|
| *Cucurbita moschata* (Outgroup) | 0 | 5 | 0.077 (0.042) | 0.058 (0.019) | 0.068 (0.029) | -0.018 (0.017) |
| Jiquipilas, Chiapas (W) | 1 | 8 | 0.091 (0.039) | 0.073 (0.020) | 0.082 (0.027) | -0.020 (0.017) |
| Ometepec, Guerrero (W) | 2 | 5 | 0.089 (0.038) | 0.076 (0.021) | 0.088 (0.030) | 0.001 (0.027) |
| Puerto Escondido, Oaxaca (W) | 3 | 14 | 0.094 (0.031) | 0.079 (0.018) | 0.085 (0.022) | -0.021 (0.017) |
| Jalisco (W) | 4 | 17 | 0.115 (0.033) | 0.099 (0.020) | 0.106 (0.024) | -0.020 (0.020) |
| Tlapehuala, Guerrero (D) | 5 | 10 | 0.098 (0.029) | 0.086 (0.018) | 0.093 (0.022) | -0.010 (0.023) |
| Jalisco (D) | 6 | 13 | 0.099 (0.025) | 0.090 (0.017) | 0.096 (0.020) | -0.006 (0.022) |
| Badiraguato, Sinaloa (D) | 7 | 1 | 0.095 (0.086) | 0.047 (0.021) | 0.095 (0.086) | 0.000 (0.000) |
| Matlalapa, Guerrero (D) | 8 | 7 | 0.092 (0.029) | 0.081 (0.018) | 0.090 (0.023) | -0.006 (0.017) |
| Sahuayo, Michoacán (D) | 9 | 10 | 0.101 (0.027) | 0.091 (0.018) | 0.097 (0.021) | -0.008 (0.022) |
| Salamanca, Guanajuato (D) | 10 | 1 | 0.087 (0.079) | 0.043 (0.020) | 0.087 (0.079) | 0.000 (0.000) |
| Tepic, Nayarit (D) | 11 | 4 | 0.089 (0.044) | 0.067 (0.020) | 0.083 (0.034) | -0.012 (0.012) |
| El Platanar, Sinaloa (feral) | 12 | 4 | 0.107 (0.062) | 0.074 (0.024) | 0.097 (0.046) | -0.019 (0.018) |
| Culiacán, Sinaloa (feral) | 13 | 3 | 0.099 (0.064) | 0.066 (0.023) | 0.094 (0.053) | -0.010 (0.014) |
| Choix, Sinaloa (D) | 14 | 3 | 0.101 (0.060) | 0.069 (0.023) | 0.096 (0.050) | -0.008 (0.014) |
| Yecora, Sonora (D) | 15 | 6 | 0.104 (0.040) | 0.081 (0.020) | 0.092 (0.027) | -0.025 (0.013) |
| Onavas, Sonora (feral) | 16 | 7 | 0.101 (0.044) | 0.077 (0.022) | 0.088 (0.029) | -0.026 (0.015) |
| Durango, Durango (D) | 17 | 1 | 0.076 (0.071) | 0.038 (0.017) | 0.076 (0.071) | 0.000 (0.000) |
| Tehuantepec, Oaxaca (D) | 18 | 7 | 0.085 (0.032) | 0.071 (0.018) | 0.079 (0.023) | -0.014 (0.014) |
| Onavas, Sonora (D) | 19 | 6 | 0.089 (0.040) | 0.067 (0.018) | 0.078 (0.026) | -0.025 (0.013) |
| Tihuatlán, Veracruz (D) | 20 | 5 | 0.088 (0.041) | 0.069 (0.020) | 0.082 (0.030) | -0.011 (0.018) |
| Palenque, Chiapas (D) | 21 | 5 | 0.089 (0.035) | 0.072 (0.019) | 0.083 (0.026) | -0.013 (0.015) |
| Tanquián, San Luis Potosí (D) | 22 | 6 | 0.090 (0.033) | 0.074 (0.019) | 0.084 (0.025) | -0.013 (0.013) |
| Champotón, Campeche (D) | 23 | 5 | 0.096 (0.039) | 0.076 (0.020) | 0.090 (0.030) | -0.011 (0.016) |
| Mixtepec, Oaxaca (D) | 24 | 6 | 0.101 (0.038) | 0.081 (0.020) | 0.092 (0.027) | -0.020 (0.016) |
| Ek Balam, Yucatan (D) | 25 | 6 | 0.098 (0.041) | 0.075 (0.020) | 0.086 (0.027) | -0.026 (0.014) |
| Chan Santa Cruz, Quintana Roo (D) | 26 | 7 | 0.089 (0.032) | 0.073 (0.018) | 0.081 (0.022) | -0.018 (0.016) |

(*N* = sample size, $H_O$ = observed heterozygosity, $H_E$ = expected heterozygosity, $\pi$ = nucleotide diversity, $F_{IS}$ = inbreeding coefficient, Var = variance) (within population names: W = wild, D = domesticated)

**Table S6.** Results of ABBA-BABA test to detect introgression using 11,498,421 variants between *C. argyrosperma* subsp. *sororia* (P1), *C. argyrosperma* subp. *argyrosperma* (P2) and *C. moschata* (P3), while using *C. okeechobeensis* subsp. *martinezii* as an outgroup.

| P1 | P2 | P3 | AABB sites | ABBA sites | BABA sites | *D*-statistic | *p*-value | $f_G$ |
|---|---|---|---|---|---|---|---|---|
| *C. argyrosperma* subsp. *sororia* | *C. argyrosperma* subsp. *argyrosperma* | *C. moschata* | 652029 | 90253.8 | 81267 | 0.0523945 | 0.001414 | 0.0106867 |

**Table S7.** Candidate genes containing at least one outlier SNP (predicted by either BayeScEnv or PCAdapt) within their inner structure (introns, exons, UTRs). The direction of selection was inferred according to the ancestral and derived allelic state for each outlier SNP. (AED = Annotation Edit Distance; GO ID = Gene Ontology ID)

| Direction of selection | Gene ID (domesticated genome) | Gene ID (wild genome) | Chromosome location | Functional annotation against SwissProt | AED | GO ID |
|---|---|---|---|---|---|---|
| domesticated populations | Carg02490 | Csor.00g220720 | Chr02 | Similar to *FZR1* Protein FIZZY-RELATED 1 (*Arabidopsis thaliana*) | 0.17 | NA |
| domesticated populations | Carg02896 | Csor.00g176320 | Chr09 | Similar to *AKT1* Potassium channel *AKT1* (*Arabidopsis thaliana*) | 0.14 | GO:0005216, GO:0006811, GO:0016020, GO:0055085 |
| domesticated populations | Carg04403 | Csor.00g121120 | Chr04 | Similar to *P4H7* Probable prolyl 4-hydroxylase 7 (*Arabidopsis thaliana*) | 0.15 | GO:0016491, GO:0055114 |
| domesticated populations | Carg04908 | Csor.00g009170 | Chr08 | Similar to *RPS15AE* 40S ribosomal protein S15a-5 (*Arabidopsis thaliana*) | 0.11 | GO:0003735, GO:0005840, GO:0006412 |
| domesticated populations | Carg05198 | Csor.00g110710 | Chr07 | Similar to *MYB44* Transcription factor *MYB44* (*Arabidopsis thaliana*) | 0.01 | GO:0003677 |
| domesticated populations | Carg06661 | Csor.00g045750 | Chr18 | Protein of unknown function | 0.15 | NA |
| domesticated populations | Carg07327 | Csor.00g196260 | Chr01 | Similar to *dusA* tRNA-dihydrouridine(20/20a) synthase (*Vibrio vulnificus* (strain CMCP6)) | 0.27 | GO:0008033, GO:0017150, GO:0050660, GO:0055114 |

| | | | | | | |
|---|---|---|---|---|---|---|
| domesticated populations | Carg09511 | Csor.00g083590 | Chr17 | Similar to *IAA27* Auxin-responsive protein *IAA27* (*Arabidopsis thaliana*) | 0.14 | NA |
| domesticated populations | Carg10909 | Csor.00g085670 | Chr05 | Protein of unknown function | 0.04 | NA |
| domesticated populations | Carg12845 | Csor.00g037030 | Chr09 | Similar to *GDPDL4* Glycerophosphodiester phosphodiesterase *GDPDL4* (*Arabidopsis thaliana*) | 0.1 | GO:0006629, GO:0008081 |
| domesticated populations | Carg13010 | Csor.00g214890 | Chr10 | Similar to At4g29530 Thiamine phosphate phosphatase-like protein (*Arabidopsis thaliana*) | 0.11 | GO:0016791 |
| domesticated populations | Carg13432 | Csor.00g005000 | Chr08 | Similar to *efr3b* Protein *EFR3* homolog B (*Danio rerio*) | 0.16 | NA |
| domesticated populations | Carg14212 | Csor.00g024080 | Chr04 | Similar to *SCAI* Protein *SCAI* (*Homo sapiens*) | 0.08 | GO:0003714, GO:0006351 |
| domesticated populations | Carg14512 | Csor.00g157750 | Chr18 | Similar to *MKP1* Protein-tyrosine-phosphatase *MKP1* (*Arabidopsis thaliana*) | 0.01 | GO:0008138, GO:0016311 |
| domesticated populations | Carg17189 | Csor.00g139890 | Chr01 | Similar to *FPP4* Filament-like plant protein 4 (*Arabidopsis thaliana*) | 0.11 | NA |
| domesticated populations | Carg18146 | Csor.00g156040 | Chr11 | Protein of unknown function | 0.13 | NA |

| | | | | | | |
|---|---|---|---|---|---|---|
| domesticated populations | Carg18171 | Csor.00g211980 | Chr11 | Similar to *AHA11* ATPase 11, plasma membrane-type (*Arabidopsis thaliana*) | 0.11 | GO:0008553, GO:0016021, GO:0120029 |
| domesticated populations | Carg18484 | Csor.00g111760 | Chr07 | Similar to At1g04910 Uncharacterized protein At1g04910 (*Arabidopsis thaliana*) | 0.25 | NA |
| domesticated populations | Carg18727 | Csor.00g105010 | Chr17 | Similar to serinc Probable serine incorporator (*Nematostella vectensis*) | 0.07 | GO:0016020 |
| domesticated populations | Carg18786 | Csor.00g231070 | Chr08 | Similar to *CURT1C* Protein CURVATURE THYLAKOID 1C, chloroplastic (*Arabidopsis thaliana*) | 0.26 | NA |
| domesticated populations | Carg20078 | Csor.00g227200 | Chr09 | Similar to *ABCE2* ABC transporter E family member 2 (*Arabidopsis thaliana*) | 0.12 | GO:0005524, GO:0016887 |
| domesticated populations | Carg20623 | Csor.00g016670 | Chr13 | Protein of unknown function | 0.03 | GO:0005515 |
| domesticated populations | Carg21521 | Csor.00g160720 | Chr13 | Similar to *SNI1* Negative regulator of systemic acquired resistance *SNI1* (*Arabidopsis thaliana*) | 0.19 | NA |
| domesticated populations | Carg22092 | Csor.00g304880 | Chr18 | Similar to *Arfrp1* ADP-ribosylation factor-related protein 1 (*Rattus norvegicus*) | 0.09 | GO:0005525 |
| domesticated populations | Carg22878 | Csor.00g267780 | Chr15 | Similar to *PTD* Protein PARTING DANCERS (*Arabidopsis thaliana*) | 0.32 | NA |

| | | | | | | |
|---|---|---|---|---|---|---|
| domesticated populations | Carg22996 | Csor.00g086420 | Chr05 | Similar to *FLK* Flowering locus K homology domain (*Arabidopsis thaliana*) | 0.11 | GO:0003723 |
| domesticated populations | Carg23167 | Csor.00g003720 | Chr08 | Similar to *CES101* G-type lectin S-receptor-like serine/threonine-protein kinase *CES101* (*Arabidopsis thaliana*) | 0.04 | GO:0004672, GO:0004674, GO:0005524, GO:0006468 |
| domesticated populations | Carg24693 | Csor.00g076650 | Chr01 | Similar to At1g17220 Translation initiation factor *IF-2*, chloroplastic (*Arabidopsis thaliana*) | 0.08 | GO:0003743, GO:0003924, GO:0005525, GO:0006413 |
| domesticated populations | Carg24979 | Csor.00g267730 | Chr15 | Similar to *Cag_1601* UPF0301 protein *Cag_1601* (*Chlorobium chlorochromatii* (strain CaD3)) | 0.29 | NA |
| domesticated populations | Carg25230 | Csor.00g030480 | Chr16 | Similar to *CLC-F* Chloride channel protein *CLC-f* (*Arabidopsis thaliana*) | 0.05 | GO:0005247, GO:0006821, GO:0016020, GO:0055085 |
| domesticated populations | Carg25231 | Csor.00g030490 | Chr16 | Similar to *WRKY2* Probable *WRKY* transcription factor 2 (*Arabidopsis thaliana*) | 0.03 | GO:0003700, GO:0006355, GO:0043565 |
| domesticated populations | Carg26216 | Csor.00g010650 | Chr15 | Similar to *FTSH11* ATP-dependent zinc metalloprotease *FTSH* 11, | 0.27 | GO:0004222, GO:0005524, |

| | | | | | | |
|---|---|---|---|---|---|---|
| | | | | chloroplastic/mitochondrial (*Arabidopsis thaliana*) | | GO:0006508, GO:0016020 |
| domesticated populations | Carg26378 | Csor.00g260430 | Chr11 | Similar to *DLO1* Protein *DMR6*-LIKE OXYGENASE 1 (*Arabidopsis thaliana*) | 0.16 | GO:0016491, GO:0055114 |
| domesticated populations | Carg26857 | Csor.00g260990 | Chr11 | Similar to *DDB_G0292320* Protein unc-50 homolog (*Dictyostelium discoideum*) | 0.24 | NA |
| domesticated populations | Carg27299 | Csor.00g267650 | Chr15 | Similar to *SAC1* Phosphoinositide phosphatase *SAC1* (*Arabidopsis thaliana*) | 0.18 | GO:0042578 |
| domesticated populations | Carg_TCONS_00100457 | NA | Chr11 | Long noncoding RNA | NA | NA |
| wild populations | Carg00678 | Csor.00g059490 | Chr03 | Similar to *CSI1* Protein CELLULOSE SYNTHASE INTERACTIVE 1 (*Arabidopsis thaliana*) | 0.06 | GO:0005515 |
| wild populations | Carg00749 | Csor.00g058770 | Chr03 | Similar to *WDR5A* COMPASS-like H3K4 histone methylase component *WDR5A* (*Arabidopsis thaliana*) | 0.01 | GO:0005515 |
| wild populations | Carg00754 | Csor.00g058710 | Chr03 | Similar to *TUBA* Tubulin alpha chain (*Prunus dulcis*) | 0.07 | GO:0003924 |
| wild populations | Carg00755 | NA | Chr03 | Protein of unknown function | 0.16 | NA |
| wild populations | Carg00942 | Csor.00g292650 | Chr06 | Similar to *CTN* Cactin (*Arabidopsis thaliana*) | 0.14 | GO:0005515 |

| | | | | | | |
|---|---|---|---|---|---|---|
| wild populations | Carg01177 | Csor.00g247900 | Chr06 | Protein of unknown function | 0.13 | GO:0071816 |
| wild populations | Carg01302 | Csor.00g249070 | Chr06 | Similar to *SSL10* Protein STRICTOSIDINE SYNTHASE-LIKE 10 (*Arabidopsis thaliana*) | 0.13 | GO:0009058, GO:0016844 |
| wild populations | Carg01309 | Csor.00g249150 | Chr06 | Similar to *ARF1* Auxin response factor 1 (*Arabidopsis thaliana*) | 0.15 | NA |
| wild populations | Carg01823 | Csor.00g164870 | Chr04 | Similar to *PP2AA2* Serine/threonine-protein phosphatase 2A 65 kDa regulatory subunit A beta isoform (*Arabidopsis thaliana*) | 0.19 | GO:0005515 |
| wild populations | Carg02429 | Csor.00g032070 | Chr15 | Protein of unknown function | 0.11 | NA |
| wild populations | Carg02612 | Csor.00g221980 | Chr02 | Similar to *TIC100* Protein *TIC* 100 (*Arabidopsis thaliana*) | 0.21 | NA |
| wild populations | Carg02996 | NA | Chr09 | Similar to At5g49980 Transport inhibitor response 1-like protein (*Arabidopsis thaliana*) | 0.02 | GO:0005515 |
| wild populations | Carg03224 | Csor.00g101370 | Chr20 | Similar to *DDB_G0284757* OTU domain-containing protein *DDB_G0284757* (*Dictyostelium discoideum*) | 0.31 | NA |

| | | | | | | |
|---|---|---|---|---|---|---|
| wild populations | Carg03669 | NA | Chr14 | Similar to At3g10130 Heme-binding-like protein At3g10130, chloroplastic (*Arabidopsis thaliana*) | 0.2 | NA |
| wild populations | Carg03798 | Csor.00g009600 | Chr08 | Similar to At1g09760 U2 small nuclear ribonucleoprotein A' (*Arabidopsis thaliana*) | 0.06 | NA |
| wild populations | Carg04098 | Csor.00g278800 | Chr19 | Similar to *GNTI* Alpha-1,3-mannosyl-glycoprotein 2-beta-N-acetylglucosaminyltransferase (*Arabidopsis thaliana*) | 0.23 | GO:0006486, GO:0008375 |
| wild populations | Carg04189 | Csor.00g196640 | Chr01 | Similar to *RAD23B* Ubiquitin receptor *RAD23b* (*Arabidopsis thaliana*) | 0.16 | GO:0003684, GO:0005515, GO:0005634, GO:0006289, GO:0043161 |
| wild populations | Carg04587 | Csor.00g123000 | Chr04 | Similar to At3g53190 Probable pectate lyase 12 (*Arabidopsis thaliana*) | 0.06 | NA |
| wild populations | Carg04919 | Csor.00g009070 | Chr08 | Similar to *U2AF65B* Splicing factor U2af large subunit B (*Nicotiana plumbaginifolia*) | 0.3 | GO:0003676, GO:0003723, GO:0005634, GO:0006397 |

| | | | | | | |
|---|---|---|---|---|---|---|
| wild populations | Carg04921 | Csor.00g009020 | Chr08 | Similar to *ATX2* Histone-lysine N-methyltransferase *ATX2* (*Arabidopsis thaliana*) | 0.23 | GO:0005515, GO:0005634 |
| wild populations | Carg04961 | Csor.00g008680 | Chr08 | Similar to *CRK3 CDPK*-related kinase 3 (*Arabidopsis thaliana*) | 0.18 | GO:0004672, GO:0005524, GO:0006468 |
| wild populations | Carg04969 | Csor.00g008610 | Chr08 | Similar to *CAT9* Cationic amino acid transporter 9, chloroplastic (*Arabidopsis thaliana*) | 0.15 | GO:0016020, GO:0022857, GO:0055085 |
| wild populations | Carg05727 | Csor.00g172100 | Chr16 | Similar to *KIN7G* Kinesin-like protein *KIN-7G* (*Arabidopsis thaliana*) | 0.11 | GO:0003777, GO:0005524, GO:0007018, GO:0008017 |
| wild populations | Carg06107 | Csor.00g118480 | Chr04 | Similar to *OPR1* 12-oxophytodienoate reductase 1 (*Arabidopsis thaliana*) | 0.32 | GO:0010181, GO:0016491, GO:0055114 |
| wild populations | Carg06628 | Csor.00g046050 | Chr18 | Similar to *NMT1* Phosphoethanolamine N-methyltransferase 1 (*Arabidopsis thaliana*) | 0.2 | GO:0008168 |
| wild populations | Carg06696 | Csor.00g045440 | Chr18 | Similar to At5g11010 Polynucleotide 5'-hydroxyl-kinase *NOL9* (*Arabidopsis thaliana*) | 0.19 | NA |

| | | | | | | |
|---|---|---|---|---|---|---|
| wild populations | Carg06792 | Csor.00g044470 | Chr18 | Similar to *MSP1* Protein *MSP1* (*Saccharomyces cerevisiae* (strain ATCC 204508 / S288c)) | 0.22 | GO:0005515, GO:0005524 |
| wild populations | Carg06849 | Csor.00g043850 | Chr18 | Similar to *SS3* Soluble starch synthase 3, chloroplastic/amyloplastic (*Solanum tuberosum*) | 0.11 | GO:2001070 |
| wild populations | Carg06968 | Csor.00g203640 | Chr01 | Similar to *ORRM6* Organelle *RRM* domain-containing protein 6, chloroplastic (*Arabidopsis thaliana*) | 0.18 | GO:0003676 |
| wild populations | Carg06997 | Csor.00g080170 | Chr01 | Similar to *IREH1* Probable serine/threonine protein kinase *IREH1* (*Arabidopsis thaliana*) | 0.08 | GO:0004672, GO:0005524, GO:0006468 |
| wild populations | Carg07232 | Csor.00g265760 | Chr17 | Similar to *HULK3* Protein *HUA2*-LIKE 3 (*Arabidopsis thaliana*) | 0.22 | NA |
| wild populations | Carg07327 | Csor.00g196260 | Chr01 | Similar to *dusA* tRNA-dihydrouridine(20/20a) synthase (*Vibrio vulnificus* (strain CMCP6)) | 0.27 | GO:0008033, GO:0017150, GO:0050660, GO:0055114 |
| wild populations | Carg07889 | Csor.00g113710 | Chr07 | Similar to *SFH9* Phosphatidylinositol/phosphatidylcholine transfer protein *SFH9* (*Arabidopsis thaliana*) | 0.14 | NA |

| | | | | | | |
|---|---|---|---|---|---|---|
| wild populations | Carg07954 | Csor.00g113070 | Chr07 | Similar to *BHLH121* Transcription factor *bHLH121* (*Arabidopsis thaliana*) | 0.03 | GO:0046983 |
| wild populations | Carg08549 | Csor.00g041110 | Chr02 | Similar to At4g10930 Uncharacterized protein At4g10930 (*Arabidopsis thaliana*) | 0.19 | NA |
| wild populations | Carg09452 | Csor.00g084200 | Chr17 | Similar to *ALA4* Probable phospholipid-transporting ATPase 4 (*Arabidopsis thaliana*) | 0.09 | GO:0000166, GO:0000287, GO:0005524, GO:0015914, GO:0016021, GO:0140326 |
| wild populations | Carg10718 | Csor.00g080500 | Chr01 | Similar to *FBL15* F-box/LRR-repeat protein 15 (*Arabidopsis thaliana*) | 0.16 | GO:0005515 |
| wild populations | Carg10970 | Csor.00g236380 | Chr08 | Similar to trc Serine/threonine-protein kinase tricorner (*Drosophila pseudoobscura pseudoobscura*) | 0.14 | GO:0004672, GO:0004674, GO:0005524, GO:0006468 |
| wild populations | Carg11153 | Csor.00g150050 | Chr14 | Protein of unknown function | 0.07 | NA |
| wild populations | Carg11621 | Csor.00g072250 | Chr02 | Similar to Zeaxanthin epoxidase, chloroplastic (*Prunus armeniaca*) | 0.12 | GO:0005515, GO:0009507, GO:0009540, GO:0009688, |

| | | | | | | |
|---|---|---|---|---|---|---|
| | | | | | | GO:0016020, GO:0055114, GO:0071949 |
| wild populations | Carg11936 | Csor.00g166690 | Chr05 | Similar to *PUB6* U-box domain-containing protein 6 (*Arabidopsis thaliana*) | 0.1 | GO:0004842, GO:0016567 |
| wild populations | Carg12108 | Csor.00g006010 | Chr05 | Similar to *ATAD1* ATPase family AAA domain-containing protein 1 (*Bos taurus*) | 0.17 | GO:0005524 |
| wild populations | Carg12374 | Csor.00g192590 | Chr01 | Similar to *PBL10* Probable serine/threonine-protein kinase *PBL10* (*Arabidopsis thaliana*) | 0.19 | GO:0004672, GO:0006468 |
| wild populations | Carg12525 | Csor.00g094660 | Chr14 | Similar to *EMB1444* Transcription factor *EMB1444* (*Arabidopsis thaliana*) | 0.24 | GO:0046983 |
| wild populations | Carg12589 | Csor.00g095250 | Chr14 | Similar to *AP2* Floral homeotic protein APETALA 2 (*Arabidopsis thaliana*) | 0.07 | GO:0003677, GO:0003700, GO:0006355 |
| wild populations | Carg13344 | Csor.00g042780 | Chr07 | Similar to *CSTF77* Cleavage stimulation factor subunit 77 (*Arabidopsis thaliana*) | 0.23 | GO:0005515, GO:0005634, GO:0006397 |
| wild populations | Carg14376 | Csor.00g056580 | Chr04 | Similar to *CLE10 CLAVATA3*/*ESR (CLE)*-related protein 10 (*Arabidopsis thaliana*) | 0.22 | NA |

| | | | | | | |
|---|---|---|---|---|---|---|
| wild populations | Carg14512 | Csor.00g157750 | Chr18 | Similar to *MKP1* Protein-tyrosine-phosphatase *MKP1* (*Arabidopsis thaliana*) | 0.01 | GO:0008138, GO:0016311 |
| wild populations | Carg14536 | Csor.00g207860 | Chr06 | Similar to At5g03900 Uncharacterized protein At5g03900, chloroplastic (*Arabidopsis thaliana*) | 0.18 | NA |
| wild populations | Carg14932 | Csor.00g116960 | Chr07 | Similar to *VPS54* Vacuolar protein sorting-associated protein 54, chloroplastic (*Arabidopsis thaliana*) | 0.26 | GO:0005515, GO:0008080, GO:0042147 |
| wild populations | Carg14976 | Csor.00g116510 | Chr07 | Similar to *ABCC2* ABC transporter C family member 2 (*Arabidopsis thaliana*) | 0.88 | GO:0005524, GO:0016021, GO:0016887, GO:0042626, GO:0055085 |
| wild populations | Carg15060 | Csor.00g282910 | Chr16 | Similar to *IQD1* Protein IQ-DOMAIN 1 (*Arabidopsis thaliana*) | 0.09 | GO:0005515 |
| wild populations | Carg15210 | Csor.00g217960 | Chr10 | Protein of unknown function | 0.33 | NA |
| wild populations | Carg15512 | Csor.00g251220 | Chr06 | Similar to *SEC23* Protein transport protein *SEC23* (*Ustilago maydis* (strain 521 / FGSC 9021)) | 0.14 | GO:0006886, GO:0006888, GO:0008270, GO:0030127 |

| | | | | | | |
|---|---|---|---|---|---|---|
| wild populations | Carg15691 | Csor.00g281360 | Chr19 | Similar to Os03g0733400 Zinc finger BED domain-containing protein *RICESLEEPER* 2 (*Oryza sativa* subsp. *japonica*) | 0.11 | GO:0003677, GO:0046983 |
| wild populations | Carg15904 | Csor.00g236940 | Chr08 | Similar to *SGS3* Protein SUPPRESSOR OF GENE SILENCING 3 homolog (*Oryza sativa* subsp. *indica*) | 0.13 | GO:0031047 |
| wild populations | Carg15929 | Csor.00g237260 | Chr08 | Similar to At5g19025 Uncharacterized protein At5g19025 (*Arabidopsis thaliana*) | 0.28 | NA |
| wild populations | Carg16055 | Csor.00g112640 | Chr07 | Similar to *AUL1* Auxilin-like protein 1 (*Arabidopsis thaliana*) | 0.04 | NA |
| wild populations | Carg16433 | NA | Chr12 | Similar to Gtp-bp Signal recognition particle receptor subunit alpha homolog (*Drosophila melanogaster*) | 0.08 | GO:0003924, GO:0005047, GO:0005525, GO:0005785, GO:0006614, GO:0006886 |
| wild populations | Carg17222 | Csor.00g304480 | Chr18 | Similar to *CUT1* 3-ketoacyl-CoA synthase 6 (*Arabidopsis thaliana*) | 0.02 | GO:0003824, GO:0006633, GO:0016020, GO:0016747 |

| | | | | | | |
|---|---|---|---|---|---|---|
| wild populations | Carg17814 | Csor.00g228870 | Chr10 | Protein of unknown function | 0.21 | NA |
| wild populations | Carg17827 | Csor.00g228730 | Chr10 | Similar to *BSL2* Serine/threonine-protein phosphatase *BSL2* (*Arabidopsis thaliana*) | 0.1 | GO:0004721, GO:0005515, GO:0009742, GO:0016787 |
| wild populations | Carg18944 | Csor.00g084780 | Chr17 | Similar to *ATL3* RING-H2 finger protein *ATL3* (*Arabidopsis thaliana*) | 0.01 | NA |
| wild populations | Carg19818 | Csor.00g017820 | Chr05 | Similar to *SPCC1672*.07 U3 small nucleolar RNA-associated protein 21 homolog (*Schizosaccharomyces pombe* (strain 972 / ATCC 24843)) | 0.18 | GO:0005515, GO:0006364, GO:0032040 |
| wild populations | Carg20889 | Csor.00g273290 | Chr06 | Protein of unknown function | 0.18 | NA |
| wild populations | Carg21706 | Csor.00g203020 | Chr01 | Similar to *DGK1* Diacylglycerol kinase 1 (*Arabidopsis thaliana*) | 0.06 | GO:0004143, GO:0007205, GO:0016301, GO:0035556 |
| wild populations | Carg22034 | Csor.00g277150 | Chr19 | Similar to *PDV2* Plastid division protein *PDV2* (*Arabidopsis thaliana*) | 0.12 | NA |
| wild populations | Carg22232 | Csor.00g242160 | Chr16 | Similar to *POP1* Ribonucleases P/MRP protein subunit *POP1* (*Homo sapiens*) | 0.1 | NA |

| | | | | | | |
|---|---|---|---|---|---|---|
| wild populations | Carg23235 | Csor.00g188450 | Chr16 | Similar to Lon protease homolog 2, peroxisomal (*Spinacia oleracea*) | 0.12 | GO:0004176, GO:0004252, GO:0005524, GO:0006508 |
| wild populations | Carg23389 | Csor.00g206240 | Chr06 | Similar to *nop12* Nucleolar protein 12 (*Schizosaccharomyces pombe* (strain 972 / ATCC 24843)) | 0.08 | GO:0003676 |
| wild populations | Carg23772 | Csor.00g220140 | Chr04 | Similar to *SAC3A SAC3* family protein A (*Arabidopsis thaliana*) | 0.14 | NA |
| wild populations | Carg23802 | NA | Chr14 | Similar to At1g06840 Probable LRR receptor-like serine/threonine-protein kinase At1g06840 (*Arabidopsis thaliana*) | 0.19 | GO:0005515 |
| wild populations | Carg24347 | Csor.00g079020 | Chr07 | Similar to *ITN1* Ankyrin repeat-containing protein *ITN1* (*Arabidopsis thaliana*) | 0.03 | NA |
| wild populations | Carg24812 | NA | Chr16 | Similar to *PBL23* Probable serine/threonine-protein kinase *PBL23* (*Arabidopsis thaliana*) | 0.07 | GO:0004672, GO:0005524, GO:0006468 |
| wild populations | Carg25109 | NA | Chr13 | Protein of unknown function | 0.23 | NA |
| wild populations | Carg25229 | Csor.00g030470 | Chr16 | Protein of unknown function | 0.05 | NA |

| | | | | | | |
|---|---|---|---|---|---|---|
| wild populations | Carg25337 | Csor.00g070030 | Chr19 | Similar to Glycerol-3-phosphate acyltransferase, chloroplastic (*Cucumis sativus*) | 0.17 | GO:0004366, GO:0006650, GO:0016746 |
| wild populations | Carg25546 | Csor.00g002770 | Chr17 | Similar to *TMKL1* Putative kinase-like protein *TMKL1* (*Arabidopsis thaliana*) | 0.3 | GO:0004672, GO:0005515, GO:0006468 |
| wild populations | Carg25626 | Csor.00g028980 | Chr01 | Similar to *ASP3* Aspartate aminotransferase 3, chloroplastic (*Arabidopsis thaliana*) | 0.14 | GO:0009058, GO:0030170 |
| wild populations | Carg25639 | Csor.00g029150 | Chr01 | Similar to *OVA7* Serine--tRNA ligase, chloroplastic/mitochondrial (*Arabidopsis thaliana*) | 0.12 | GO:0000166, GO:0004812, GO:0004828, GO:0005524, GO:0006418, GO:0006434 |
| wild populations | Carg26784 | NA | Chr15 | Protein of unknown function | 0.02 | NA |
| wild populations | Carg26826 | Csor.00g030720 | Chr16 | Similar to *PA200* Proteasome activator subunit 4 (*Arabidopsis thaliana*) | 0.14 | NA |
| wild populations | Carg26907 | Csor.00g220120 | Chr04 | Similar to *maea* Macrophage erythroblast attacher (*Danio rerio*) | 0.14 | NA |

| | | | | | | |
|---|---|---|---|---|---|---|
| wild populations | Carg27113 | Csor.00g014920 | Chr19 | Similar to *VPS13C* Vacuolar protein sorting-associated protein 13C (*Homo sapiens*) | 0.11 | NA |
| wild populations | Carg27622 | Csor.00g103900 | Chr20 | Similar to *SUMO2* Small ubiquitin-related modifier 2 (*Arabidopsis thaliana*) | 0.32 | NA |
| wild populations | Carg_TCONS_00026631 | NA | Chr13 | Long noncoding RNA | NA | NA |
| wild populations | Carg_TCONS_00098388 | NA | Chr12 | Long noncoding RNA (pseudogene homolog of trafficking protein particle complex subunit 11) | NA | NA |
| wild populations | Carg_TCONS_00098395 | NA | Chr12 | Long noncoding RNA (pseudogene homolog of trafficking protein particle complex subunit 11) | NA | NA |
| wild populations | Carg_TCONS_00100643 | NA | Chr16 | Long noncoding RNA | NA | NA |
| ABBA sites | Carg04098 | Csor.00g278800 | Chr19 | Similar to *GNTI* Alpha-1,3-mannosyl-glycoprotein 2-beta-N-acetylglucosaminyltransferase (*Arabidopsis thaliana*) | 0.23 | GO:0006486, GO:0008375 |
| ABBA sites | Carg07674 | Csor.00g064770 | Chr13 | Similar to *apaG* Protein *ApaG* (*Magnetospirillum magneticum* (strain AMB-1 / ATCC 700264)) | 0.21 | GO:0005515 |

| | | | | | | |
|---|---|---|---|---|---|---|
| ABBA sites | Carg13651 | Csor.00g132700 | Chr02 | Similar to *Unc45a* Protein unc-45 homolog A (*Mus musculus*) | 0.12 | GO:0005515 |
| ABBA sites | Carg22996 | Csor.00g086420 | Chr05 | Similar to *FLK* Flowering locus K homology domain (*Arabidopsis thaliana*) | 0.11 | GO:0003723 |
| ABBA sites | Carg26784 | NA | Chr15 | Protein of unknown function | 0.02 | NA |
| ABBA sites | Carg26826 | Csor.00g030720 | Chr16 | Similar to *PA200* Proteasome activator subunit 4 (*Arabidopsis thaliana*) | 0.14 | NA |
| ABBA sites | Carg_TCONS_00015730 | NA | Chr03 | Long noncoding RNA | NA | NA |
| ABBA sites | Carg_TCONS_00016456 | NA | Chr03 | Long noncoding RNA | NA | NA |
| BABA sites | Carg02996 | NA | Chr09 | Similar to At5g49980 Transport inhibitor response 1-like protein (*Arabidopsis thaliana*) | 0.02 | GO:0005515 |
| BABA sites | Carg04520 | Csor.00g122340 | Chr04 | Similar to *RBCMT* Ribulose-1,5 bisphosphate carboxylase/oxygenase large subunit N-methyltransferase, chloroplastic (*Nicotiana tabacum*) | 0.23 | GO:0005515 |
| BABA sites | Carg13413 | Csor.00g005190 | Chr08 | Similar to Sacs Sacsin (*Mus musculus*) | 0.06 | NA |
| BABA sites | Carg20078 | Csor.00g227200 | Chr09 | Similar to *ABCE2* ABC transporter E family member 2 (*Arabidopsis thaliana*) | 0.12 | GO:0005524, GO:0016887 |
| BABA sites | Carg21397 | Csor.00g161930 | Chr13 | Similar to *AGD12* ADP-ribosylation factor GTPase-activating protein AGD12 (*Arabidopsis thaliana*) | 0.16 | GO:0005096 |

| | | | | | | |
|---|---|---|---|---|---|---|
| BABA sites | Carg22875 | Csor.00g267800 | Chr15 | Similar to *SPBC3E7*.09 Uncharacterized protein slp1 (Schizosaccharomyces pombe (strain 972 / ATCC 24843)) | 0.11 | NA |
| Unknown direction | Carg05757 | Csor.00g172420 | Chr16 | Similar to *BGAL3* Beta-galactosidase 3 (*Arabidopsis thaliana*) | 0.17 | GO:0030246 |

**Table S8.** Significantly enriched (*p*-value < 0.05) Gene Ontology terms for the 125 candidate genes. (GO ID = Gene Ontology ID)

| Biological Process | | |
|---|---|---|
| **GO ID** | **Term** | ***p*-value** |
| GO:0009688 | Abscisic acid biosynthetic process | 0.0055 |
| GO:0006434 | Seryl-tRNA aminoacylation | 0.0111 |
| GO:0071816 | Tail-anchored membrane protein insertion into ER membrane | 0.0111 |
| GO:0009742 | Brassinosteroid mediated signaling pathway | 0.0274 |
| GO:0042147 | Retrograde transport, endosome to Golgi | 0.0328 |
| GO:0043161 | Proteasome-mediated ubiquitin-dependent protein catabolic process | 0.0328 |
| GO:0007205 | Protein kinase C-activating G protein-coupled receptor signaling pathway | 0.0382 |
| **Molecular Function** | | |
| **GO ID** | **Term** | ***p*-value** |
| GO:0009540 | Zeaxanthin epoxidase [overall] activity | 0.0059 |
| GO:0004828 | Serine-tRNA ligase activity | 0.0117 |
| GO:0004366 | Glycerol-3-phosphate O-acyltransferase activity | 0.0117 |
| GO:0004176 | ATP-dependent peptidase activity | 0.0117 |
| GO:0005515 | Protein binding | 0.0148 |
| GO:0003714 | Transcription corepressor activity | 0.0233 |
| GO:0005047 | Signal recognition particle binding | 0.0291 |
| GO:0004674 | Protein serine/threonine kinase activity | 0.0324 |
| GO:0017150 | tRNA dihydrouridine synthase activity | 0.0348 |
| GO:0004143 | Diacylglycerol kinase activity | 0.0405 |
| GO:0016844 | Strictosidine synthase activity | 0.0405 |
| GO:2001070 | Starch binding | 0.0405 |
| **Cellular Component** | | |
| **GO ID** | **Term** | ***p*-value** |
| GO:0005785 | Signal recognition particle receptor complex | 0.011 |
| GO:0032040 | Small-subunit processome | 0.043 |

**Table S9.** Structural variants (SVs) with putative signals of selection.

| Direction of putative selection | Types of SVs | SV chromosome | SV position (bp) | Num. of candidate SNPs within SV | Method to detect candidate SNP |
|---|---|---|---|---|---|
| selective signals in *argyrosperma* | Copy-gain variants: 2 | Chr11 | 10306765-10325954 | 1 | PCAdapt |
| | | Chr13 | 5018400-5031207 | 1 | PCAdapt |
| | Copy-loss variants: 4 | Chr11 | 6731911-6735114 | 2 | PCAdapt |
| | | Chr16 | 81804-86522 | 2 | BayeScEnv |
| | | Chr16 | 8156199-8175733 | 1 | PCAdapt |
| | | Chr18 | 8143260-8151300 | 1 | PCAdapt |
| | Translocations: 3 | Chr14 | 9363022-9380874 | 4 | PCAdapt |
| | | Chr16 | 81556-85865 | 2 | BayeScEnv |
| | | Chr19 | 5610649-5616432 | 1 | BayeScEnv |
| | Unalignable regions: 1 | Chr13 | 3481538-3512101 | 2 | BayeScEnv |
| selective signals in *sororia* | Copy-gain variants: 1 | Chr12 | 1402937-1404616 | 1 | BayeScEnv |
| | Copy-loss variants: 5 | Chr04 | 11758235-11765862 | 1 | PCAdapt |
| | | Chr11 | 6079664-6092872 | 3 | BayeScEnv |
| | | Chr16 | 8156199-8178291 | 4 | BayeScEnv |
| | | Chr18 | 8143260-8151300 | 2 | PCAdapt |
| | | Chr20 | 8112234-8117389 | 1 | BayeScEnv |
| | Inversions: 3 | Chr07 | 4511851-4699925 | 1 | PCAdapt |
| | | Chr09 | 8452514-8462920 | 1 | PCAdapt |
| | | Chr12 | 5913285-6061418 | 1 | PCAdapt |
| | Translocations: 14 | Chr03 | 9228029-10350919 | 1 | PCAdapt |
| | | Chr05 | 9781718-9973041 | 1 | PCAdapt |
| | | Chr08 | 6565856-6810950 | 1 | PCAdapt |
| | | Chr09 | 4672489-4814550 | 1 | PCAdapt & BayeScEnv |
| | | Chr11 | 5643610-5706361 | 1 | PCAdapt |
| | | Chr11 | 7104627-7222534 | 1 | PCAdapt |
| | | Chr12 | 1401624-1419318 | 1 | BayeScEnv |
| | | Chr14 | 9363022-9380874 | 1 | PCAdapt |

| | | | | | |
|---|---|---|---|---|---|
| | | Chr14 | 12579480-12651643 | 6 | PCAdapt |
| | | Chr15 | 5271743-5318056 | 1 | BayeScEnv |
| | | Chr15 | 6571926-6591899 | 1 | PCAdapt |
| | | Chr15 | 7598929-7630690 | 1 | PCAdapt |
| | | Chr16 | 8159027-8178291 | 2 | PCAdapt |
| | | Chr19 | 3940354-3976131 | 1 | BayeScEnv |
| | Unalignable regions: 11 | Chr01 | 10912781-10929805 | 1 | PCAdapt & BayeScEnv |
| | | Chr05 | 5221727-5259838 | 1 | PCAdapt & BayeScEnv |
| | | Chr09 | 10882200-10916261 | 1 | PCAdapt |
| | | Chr11 | 8292116-8297494 | 1 | PCAdapt |
| | | Chr13 | 807870-850245 | 2 | PCAdapt |
| | | Chr13 | 3481538-3512101 | 2 | BayeScEnv |
| | | Chr14 | 8984610-9118533 | 1 | BayeScEnv |
| | | Chr15 | 6561310-6571925 | 1 | PCAdapt |
| | | Chr16 | 5604459-5613788 | 1 | PCAdapt |
| | | Chr18 | 3817767-3875500 | 1 | BayeScEnv |
| | | Chr20 | 7813844-7819199 | 1 | BayeScEnv |
| Selective signals of *argyrosperma x moschata* introgression | Copy-gain variants: 1 | Chr04 | 16412840-16519205 | 1 | PCAdapt & BayeScEnv |
| | Translocations: 1 | Chr08 | 6565856-6810950 | 1 | PCAdapt |
| | Unalignable regions: 1 | Chr20 | 7813844-7819199 | 1 | BayeScEnv |
| Selective signals with unknown direction | Copy-gain variants: 1 | Chr04 | 11343456-11346173 | 1 | PCAdapt |
| | Copy-loss variants: 2 | Chr03 | 2395365-2397235 | 2 | BayeScEnv |
| | | Chr16 | 81804-86522 | 1 | BayeScEnv |
| | Translocations: 6 | Chr03 | 1-102830 | 1 | BayeScEnv |
| | | Chr03 | 2395365-2412218 | 2 | BayeScEnv |
| | | Chr13 | 3035244-3053594 | 1 | PCAdapt |
| | | Chr16 | 81556-85865 | 1 | BayeScEnv |

| | | | | |
|---|---|---|---|---|
| | Chr18 | 6690077-6712190 | 2 | PCAdapt |
| | Chr20 | 7751549-7789394 | 3 | PCAdapt |
| | Chr09 | 7651180-7728149 | 1 | BayeScEnv |
| Unalignable regions: 3 | Chr11 | 8965560-8973537 | 2 | BayeScEnv |
| | Chr15 | 6561310-6571925 | 2 | PCAdapt |

### Supplementary Figures

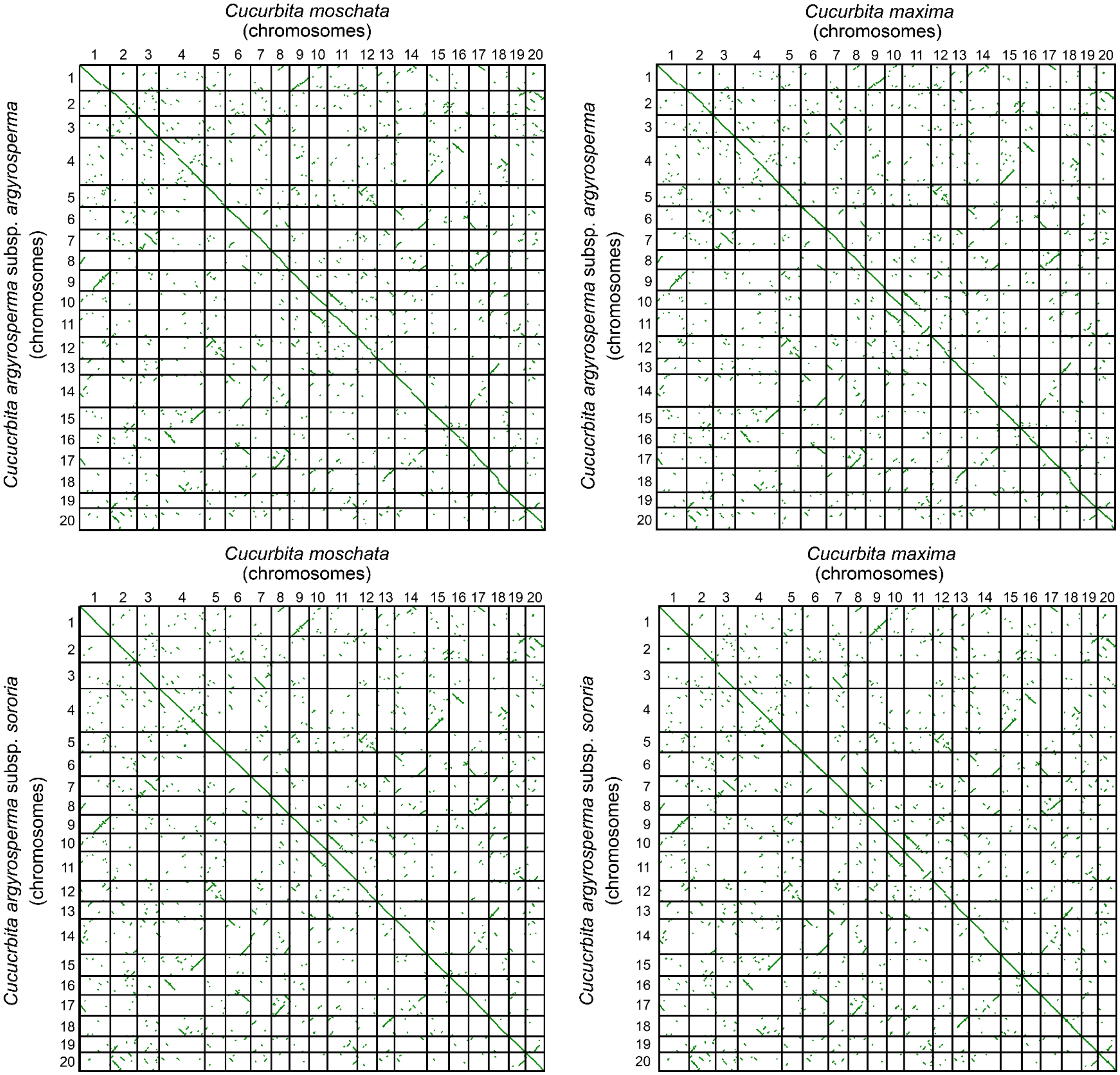


**Figure S1.** Synteny dot plots between the chromosome-level genome assemblies of *Cucurbita argyrosperma* subsp. *argyrosperma* and *C. argyrosperma* subsp. *sororia* against the reference genomes of *C. moschata* and *C. maxima* (Sun *et al*., 2017). Most of the chromosomes show chromosome-wide homoeologous pairs within the genome assembly, which have been previously attributed to a whole-genome duplication event in the *Cucurbita* genus.

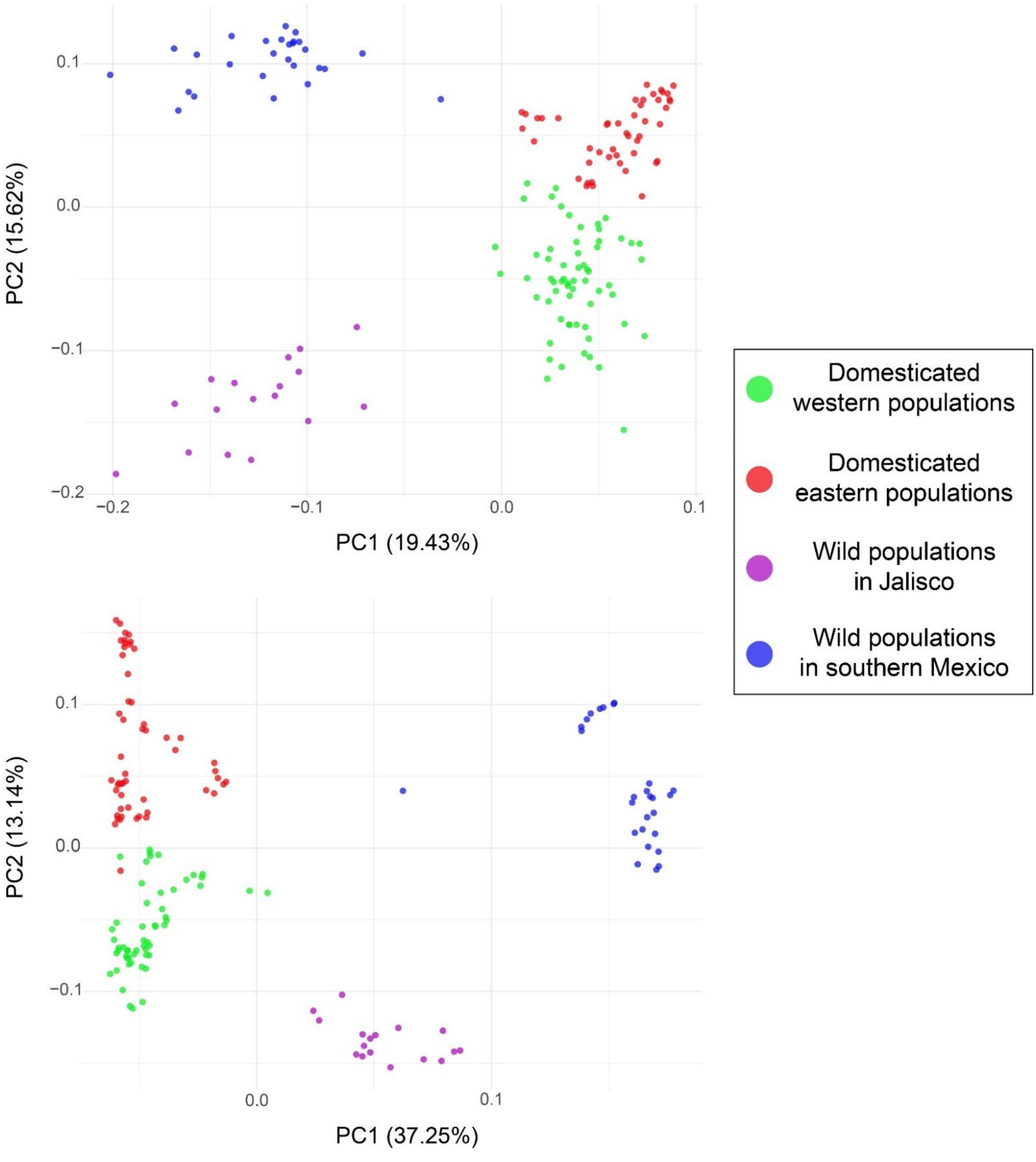


**Figure S2.** Principal Component Analysis (PCA) plots from data of *C. argyrosperma* subsp. *argyrosperma* (domesticated) and *C. argyrosperma* subsp. *sororia* (wild) populations. The first two components were plotted for (A) the SNPs dataset used for demographic analyses (2,861 SNPs,153 individuals) and for (B) the SNP dataset used for the selection scans (10,617 SNPs, 153 individuals).

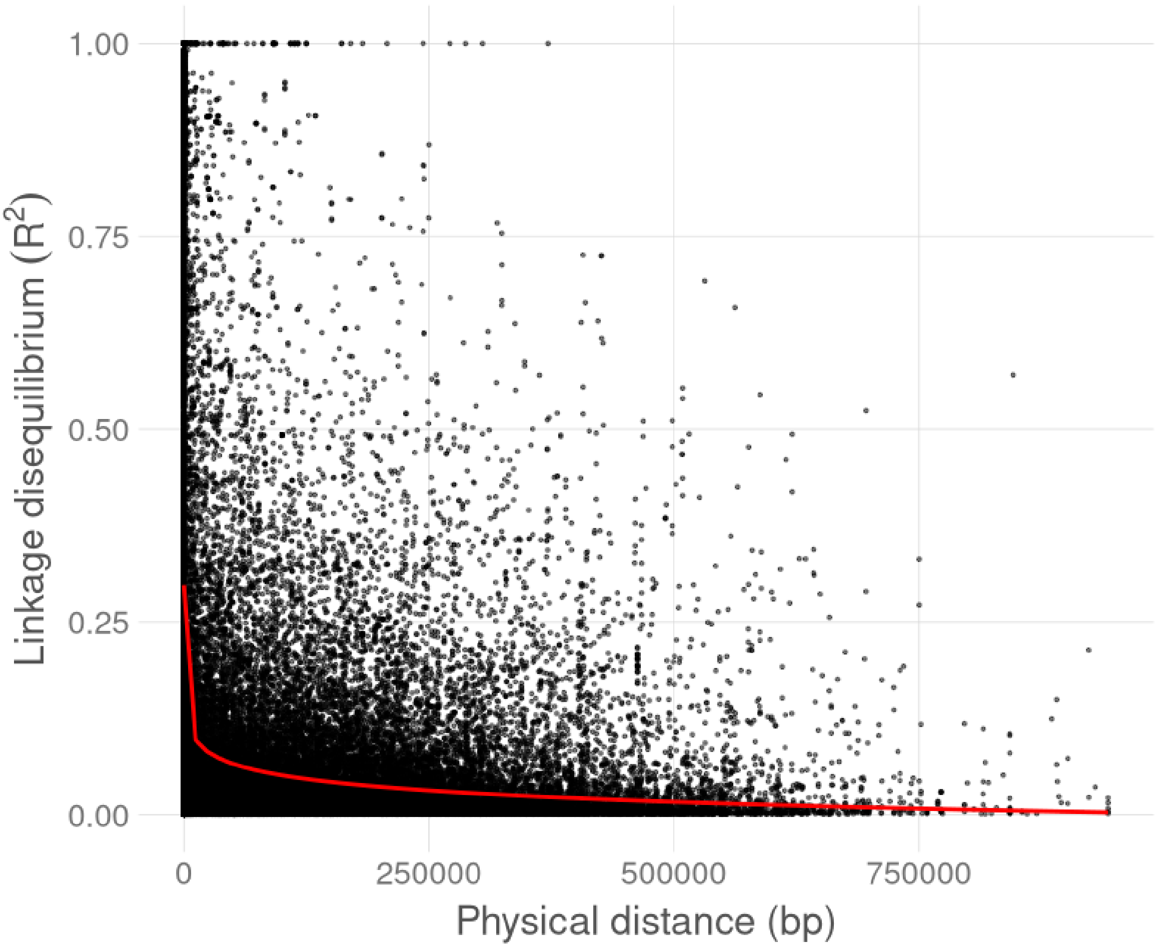


**Figure S3.** LD decay graph between the 10,617 SNPs used to perform the selective scans in *C. argyrosperma*.

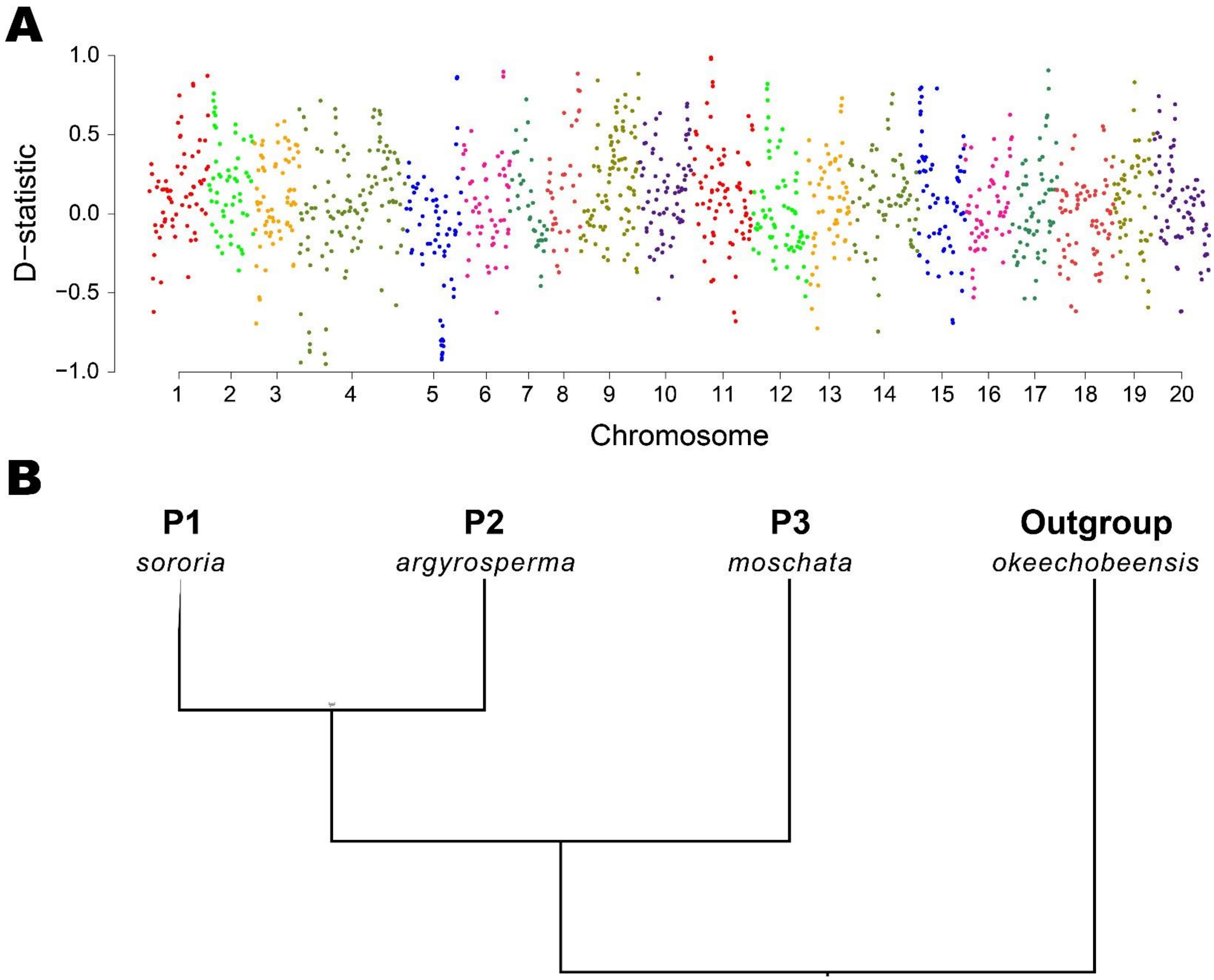


**Figure S4.** ABBA-BABA test to detect introgression using 11,498,421 variants between *C. argyrosperma* subsp. *sororia* (P1), *C. argyrosperma* subp. *argyrosperma* (P2) and *C. moschata* (P3), while using *C. okeechobeensis* subsp. *martinezii* as an outgroup. (A) Manhattan plot of the *D*-statistic throughout the genome using 500 SNP windows with a step size of 250 SNPs. (B) Phylogenetic topology used to perform the ABBA-BABA test, as reconstructed by Dsuite.
